# Aberrant formation of long-range projections across different neurodevelopmental disorders converges on molecular and cellular nexuses

**DOI:** 10.1101/2025.04.15.648981

**Authors:** Federica Furlanetto, Alejandro Segura, Mathar Kravikass, Pritha Dolai, Sarah Frank, Sören Turan, Angelica Luna Leal, Stephan Käseberg, Susann Schweiger, Chichung D. Lie, Vasily Zaburdaev, Marisa Karow, Sven Falk

## Abstract

Establishing long-range connections during human brain development is an intricate multi-step process disturbed in many neurodevelopmental disorders (NDDs). The aberrant formation of these connections is caused by mutations in a plethora of different genes with distinct molecular functions, triggering the question of whether there are common key downstream mediators at which different pathologies are converging. We employed brain organoids to model early human brain developmental aspects of Coffin-Siris-like 9, Opitz BBB/G, and Pitt-Hopkins syndromes. These NDDs are caused by mutations in *SOX11*, *MID1*, and *TCF4* respectively, and are characterized by a multitude of distinct symptoms yet share alterations in long-range projections as a common feature. Here, we uncover that mutations in all three genes phenotypically converge, showing impaired neurite extension with increased tortuosity and decreased growth speed resulting in shorter beelines. Moreover, the mutant neurites exhibit a decrease in growth persistence providing a conceptual framework explaining why long- but not short-range connections are affected. Correlating with the converging cellular phenotype, molecular characterization revealed a striking convergence on signaling pathways implicated in the interaction of neurites with their extracellular environment. *In-silico* modeling and perturbation of neurite outgrowth suggest that altered neurite-extracellular environment interactions are sufficient to recapitulate the mutant phenotypes but also facilitate the prediction of specific parameters causing disturbed neurite growth in mutant neurons.

## Introduction

In neurodevelopmental disorders (NDDs), the coordination of molecular and cellular frameworks controlling neural progenitor proliferation, neuronal differentiation and migration as well as the growth of neuronal projections is disturbed^1,2^, oftentimes leading to abnormal formation of longrange projections^3,4^. Long-range neuronal projections, connecting cortical regions with each other and with subcortical structures, include the corpus callosum, corticospinal tract, corticothalamic projections, and thalamocortical projections^1^. The proper establishment of long-range projections relies on the coordinated acquisition of neuronal identities, neuronal migration, and pathfinding of neuronal projections in the extracellular matrix (ECM)^3,5–8^. Interestingly, albeit being caused by mutations in distinct genes causing NDDs with various distinct symptoms, malformation of longrange projections is a widely observed shared symptom among NDDs^9^. Yet, how mutations in different genes coding for proteins with different molecular functions can induce a shared phenotype in humans is not well understood. With the advent of brain organoids derived from human pluripotent stem cells (hPSCs) and endowed with (patient-) specific mutations, we have now the technical possibility to tackle such questions^10–13^.

In our study, we set out to perform a comparative analysis of phenotypes underlying different NDDs that share aberrations in long-range neuronal projections as a common feature. The rare autosomal dominant disorder Coffin-Siris-like syndrome 9 (CSS9; OMIM #615866) is caused by heterozygous mutations in the gene *SOX11* (SRY-Box 11)^14^. Key symptoms of CSS9 include developmental delay, intellectual disability, microcephaly, skeletal abnormalities, and various congenital anomalies affecting the cardiac, gastrointestinal, genitourinary, and central nervous system. S*OX11* encodes a transcription factor (TF) essential for early neurodevelopment and neural differentiation^14–16^. SOX proteins contain an SRY box, a high mobility group (HMG) DNA-binding domain, allowing them to act as transcriptional activators or repressors in gene regulation. In mice, *Sox11* deletion impairs cortical neurogenesis due to reduced proliferation and abnormal differentiation of neuronal progenitor cells^17^. Sox11 physically interacts with Tcf4 (transcription factor 4) with double haploinsufficient mice showing stronger phenotypes in the truncation of the corpus callosum and the formation of the anterior commissure highlighting additive activities of Sox11 and Tcf4 in the formation of long-range connections^18^. Heterozygous mutations in *TCF4* cause the rare genetic disorder Pitt-Hopkins syndrome (PTHS; OMIM #610954). This syndrome is marked by a spectrum of developmental and physical challenges, including moderate-to-severe intellectual disability, delayed motor skills, and significant speech impairment^19,20^, resulting from *TCF4* haploinsufficiency. MRI imaging data often reveal abnormalities like white matter hyperintensity in the temporal poles, reduced hippocampal size, and mild frontal lobe hypoplasia, while EEGs may show atypical brain wave patterns, especially in patients with seizures^21^. TCF4, also known as “ITF-2” or “SEF2,” belongs to the class I basic helix-loop-helix (bHLH) family and is essential for proper cortex development^22,23^, neuronal migration^24,25^, and oligodendrogenesis^26–28^. In addition to these two TF causing NDDs we focused on the microtubule associated ubiquitin ligase MID1. Mutations in the X-linked *MID1* (midline-1) gene cause Opitz BBB/G syndrome (OS; OMIM #300000)^29^. A wide range of malformations are observed in patients with OS, affecting multiple structures during embryonic development. Common craniofacial defects include hypertelorism, often associated with a broad and flattened nasal bridge, as well as a bifid nasal tip, prominent forehead, microcephaly, cleft lip, and cleft palate^30^. Brain anomalies include hypoplasia of the cerebellar vermis, agenesis or hypoplasia of the corpus callosum, enlarged magna cisterna, and Dandy-Walker malformation^31^. MID1 functions as an E3 ubiquitin ligase, tagging target proteins with ubiquitin to regulate their degradation and function. This protein associates with microtubules and forms a multiprotein complex with protein phosphatase 2A (PP2A)^32–34^.

While the three different NDDs CSS9, OS, and PTHS include a wide spectrum of clinical symptoms, they share aberrations in the formation of long-range neuronal projections. To study these NDDs we generated brain organoids derived from isogenic pairs of human pluripotent stem cells (hPSCs) carrying loss of function mutations in either the *SOX11*, the *TCF4* or the *MID1* gene and conducted comparative molecular and cellular phenotyping of the developing brain-like tissue.

## Results

### Transiently altered neurogenesis in SOX11, TCF4, and MID1 variant brain organoids

We employed three sets of hPSCs, in which the disease-associated mutations were introduced via CRISPR/Cas9 genome editing, resulting in distinct lines, each paired with a corresponding isogenic wild-type clone. SOX11 HET carries a heterozygous mutation in the *SOX11* gene and is derived from the wildtype H9 human embryonic stem cell line which we referred to as SOX11 WT. TCF4 HET, is derived from the wildtype Hues6 human embryonic stem cell line (referred to as TCF4 WT) and harbors a heterozygous mutation resulting in TCF4 protein reduction (Figures S1B, C). In both mutant lines, SOX11 HET and TCF4 HET the mutations led to frameshifts that in case of translation would truncate the protein. MID1 KO is based on a male human induced pluripotent stem cell line (hiPSC) referred to as MID1WT in which the X-chromosomal MID1 gene is deleted hemizygously^35^ (Figures 1A-C, Figure S1A, B). Importantly, all lines retained expression of pluripotency markers, indicating that these mutations did not affect their stemness properties (Figures S1D-F). Brain organoids were generated from each hPSC line, and cellular phenotyping was performed at 30 and 60 days post aggregation of the hPSCs (Figure 1D). Using an automated image analysis pipeline, we quantified the abundance of the progenitor cell marker SOX2 and the neuronal marker MAP2. Significant reduction of the MAP2 positive area was observed in SOX11 HET (Figures 1E, F) and MID1 KO (Figures 1G, H) organoids, indicating a decrease in neurons compared to controls, while TCF4 HET organoids did not show this phenotype (Figures 1I, J). Concomitantly, an increase in the fraction of SOX2 positive area was noted in SOX11 HET and MID1 KO organoids, but not in TCF4 HET (Figure S2A). Further analysis of the tissue architecture of the d30 organoids with a special focus on ventricular zone-like structures (VZLS) revealed a shared increase in total VZLS area in both SOX11 HET and MID1 KO organoids (Figure S1B). However, we found further phenotypic differences between the two lines: MID1 KO organoids showed increased VZLS thickness (Figure S1C), while SOX11 HET organoids exhibited a significant elongation at the apical length of the VZLS (Figure S1D). This data suggests distinct modes of progenitor expansion, with a radial increase in MID1 KO organoids and a lateral increase of the VZLS in SOX11 HET organoids. Supporting this observation, a higher fraction of non-apical divisions was found in MID1 KO organoids, as indicated by the distribution of mitotic cells marked by phospho-Histone H3 (Figures S2E, F). In TCF4 HET organoids, none of these parameters were affected, though the atypical presence of cells expressing the neuronal marker DCX within the VZLS was observed (Figure S1G). By day 60, the structural differences between WT and mutant cells were no longer statistically significant across conditions (Figures 1K-P). In sum, the comparative tissue phenotyping of the organoids revealed that SOX11 and MID1 variants had a transient impact on the production of neurons, which was compensated for by day 60. Thus, we observed no long-lasting overall deficit in the formation of neurons across conditions.

**Figure 1.**
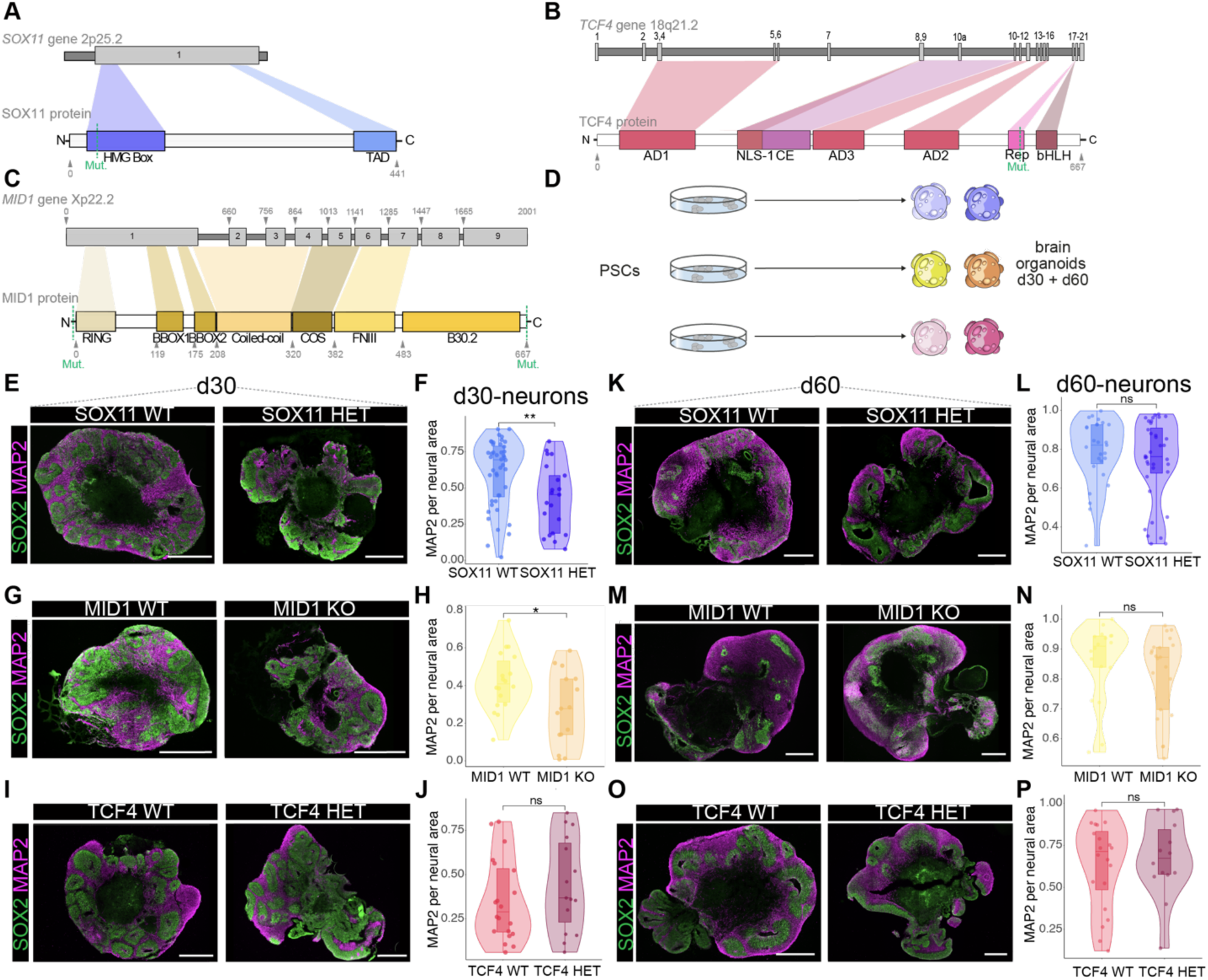
Transiently altered neurogenesis in SOX11, TCF4, and MID1 variant brain organoids. **A,** Gene and protein structure indicating the mutations in SOX11 HET, **B,** TCF4 HET, and **C,** MID1 KO PSC lines. **D,** Experimental scheme. **E, G, I,** Images showing immunohistochemical stainings of brain organoid slices (d30) stained for SOX2 and MAP2. Scale bars = 500 µm. **F, H, J,** Quantification of MAP2 signal per neural area (SOX2 and/or MAP2 signal) showing significantly decreased neurons in SOX11 HET and MID1 KO as shown by violin, box, and jitter plots. For **F,** SOX11 WT: *n* = 44; SOX11 HET: n = 21, exact *P* value 0.0054. For **H,** MID1 WT: *n* = 18; MID1 KO: n = 15, exact *P* value 0.024. For **J,** TCF4 WT: *n* = 20; TCF4 HET: n = 15, exact *P* value 0.28. **K, M, O,** Images showing brain organoid slices (d60) stained for SOX2 and MAP2. Scale bars = 500 µm. **L, N, P,** Quantification of MAP2 signal per neural area (SOX2 and/or MAP2 signal) showing no significant differences in neurons across conditions as shown by violin, box plots and jitter. For **L,** SOX11 WT: *n* = 11; SOX11 HET: n = 11, exact *P* value 0.32. For **N,** MID1 WT: *n* = 18; MID1 KO: n = 21, exact *P* value 0.33. For **P,** TCF4 WT: *n* = 20; TCF4 HET: n = 14, exact *P* value 0.61. **F, H, J, L, N, P,** \*\*\**P*<0.001, \*\**P*<0.01, \**P*<0.5, ns = non-significant; dots represent individual organoids; one-way anova with tukey posthoc test was performed.

### Live imaging reveals impaired neurite growth

Next, we wondered whether the neurons produced would differ in cellular features required for the formation of long-range projections. Live imaging was conducted to record neurite outgrowth by slicing day 60 organoids, plating on matrigel coated wells, and labeling of the microtubule cytoskeleton by SiR-tubulin, allowing real-time tracking of neurite growth over 72 hours with images taken every 20 minutes (Figure S2H). The tips of individual growing neurites were tracked as exemplified (Figures 2A-C; supplementary movies M1-M6). By inspecting the movies of the growing neurites, we have noticed that, in fact, their tips follow a trajectory that is reminiscent of the trajectory of a persistent random walk, while the shape of the grown neurite itself is similar to that of a semi-flexible polymer. Interestingly, the statistical characteristics that are commonly used to describe random walk trajectories and polymer shapes are essentially the same. Thus, we decided to use those quantifiers for the description of the neurite growth in our experiments. To this end, we analyzed the trajectories traveled by the tips of the growing neurites obtained for different conditions. This provided us with a set of data describing the two-dimensional coordinates of the growing neurite tip at discrete time points. We first quantified the instantaneous growth speed of neurites for varying time increments averaged along the trajectory and for all trajectories of the same condition. Speed calculated for small time increments is generally higher reflecting the effect of rapid growth fluctuations, while for larger increments, high-frequency fluctuations are averaged out providing the characteristic speed of neurite growth (Figure 2D). Strikingly, this analysis revealed that neurites from all conditions SOX11 HET, MID1 KO, and TCF4 HET showed a reduced speed of growth as compared to their WT counterparts (Figure 2D). We next evaluated the length of the path *L* traveled by the tip as a function of time (Figure S2I). We observed that the track length grows approximately linearly with time indicating overall constant growth speed (slope of the curves). Consistently with our previous result, we see that for the same duration, track length and growth speed are smaller in the case of SOX11 HET, MID1 KO, and TCF4 HET neurites (Figure S2I). To assess how far the neurite tips travel we quantified the end-to-end distance *d* from the starting point to the current position of the tip as a function of time. Fitting of the data to the form *d* = *a* + *b*√𝑡 (dashed lines in Figure 2E) highlights that the experimental data closely follow the prediction from the theory of random walks, where the end-to-end distance scales proportionally to the square root of time^36^. Again, we see that across conditions SOX11 HET, MID1 KO, and TCF4 HET neurites travel shorter distances. While the length of the track grows linearly in time, the neurite tip traces a complex path with continuous changes in directionality of growth. To quantify this complexity, we calculate the tortuosity *T*, which is the ratio of the path length *L* to the end-to-end distance *T*=*L*/d, with lower values for T indicating a more directed growth (*T*=1 corresponds to the growth along the straight path). Interestingly, again all mutants show an increased tortuosity (Figure 2F) highlighting that the directionality of their growth is reduced and more stochastic indicating their exploratory behavior is changed. To directly quantify the exploratory behavior of a random-walk-like trajectory we calculated the Mean Squared Displacement (MSD) as a function of time. For a normal diffusion process, MSD scales linearly with time, where the prefactor provides the estimate of the diffusion constant^37^. In the experimental data this linear behavior is only established at large enough times. For shorter times, the behavior is superdiffusive (faster than diffusion) (Figure 2G). This indicates that for short times, the direction of neurite growth remains correlated which leads to a persistent (directed) growth. However, this persistence in growth directionality is gradually lost after a certain time and the behavior of the tip becomes diffusive growing in more random directions ‘forgetting’ the initial growth direction (Figure 2G). To directly test for correlation in neurite growth direction we calculated the velocity auto-correlation function (VACF) which quantifies correlations in neurite growth velocities separated by a certain lag time. As a result, we obtain a function that decays with the lag time and exhibits two distinct regimes (see Figure 2F) of rapid decay for very short times (of the order of our temporal resolution) and a slower decay for longer times. The first regime shows the loss of correlation due to rapid velocity fluctuations, while the slower decay corresponds to the persistence phase. Such a behavior of VACF is typical for the tracking of biological movements^38^. If we re-plot the VACF in the log-linear scale (see the insets in Figure 2F), we see that it looks like a composition of two straight lines with different slopes, indicative of exponential decays. We used a simple fitting function 𝐴𝑒^−𝑡/𝜏1^ + 𝐵𝑒^−𝑡/𝜏2^ and focused our attention on the slowest time scale 𝝉_2_ that we call a persistence time. We see that in the SOX11 HET and MID1 KO, correlations decay faster and with a shorter characteristic time than in the WT conditions, and in the TCF4 HET they show the same yet not so pronounced trend (Figure 2I). Taken together, we observe that mutations in SOX11, MID1, and TCF4 lead to slower growing protrusions with lower persistence time and increased tortuosity. These phenotypes, shared among all three conditions, highlight a mechanistic convergence on a cellular level in neurite growth despite distinct underlying genetic aberrations.

**Figure 2.**
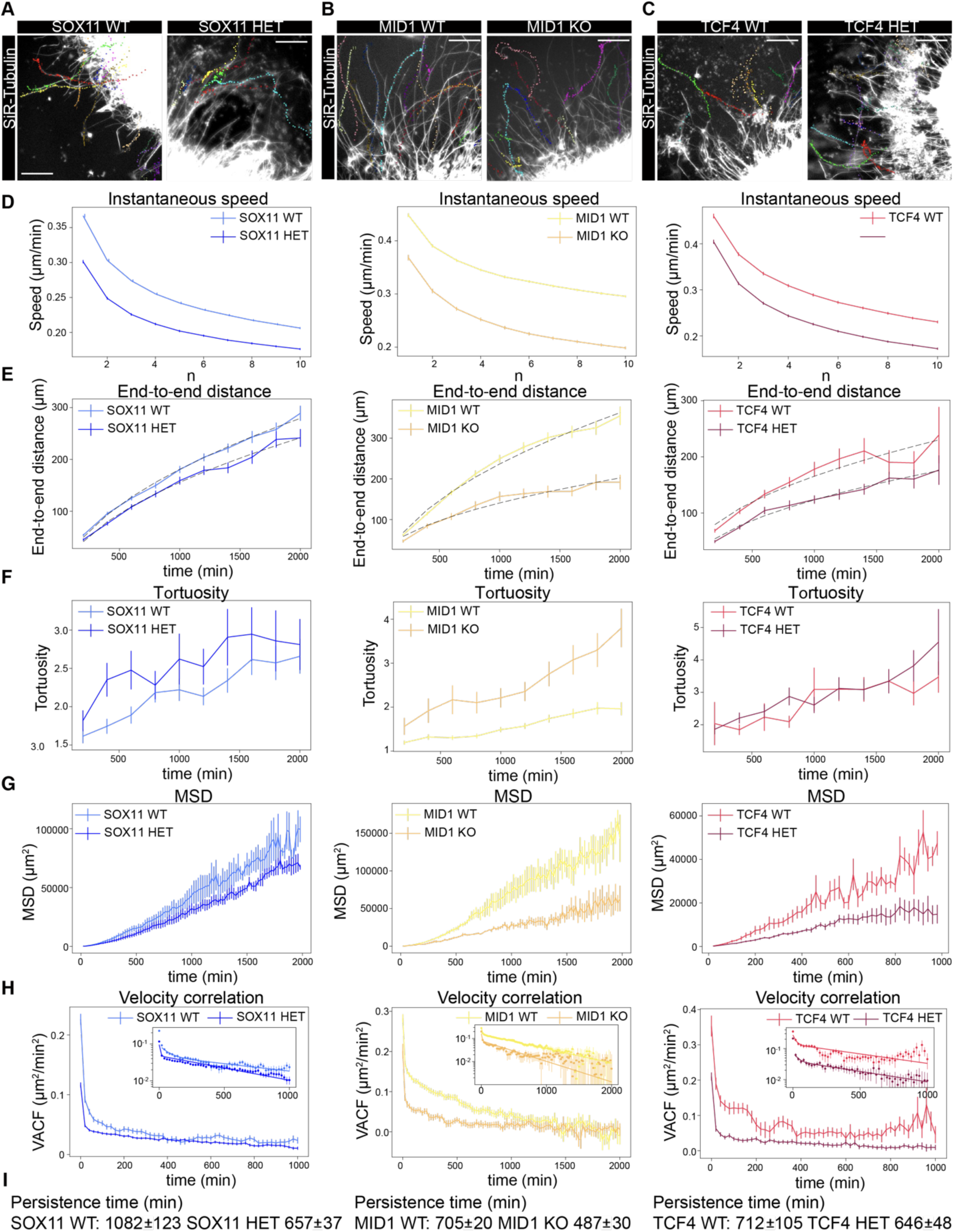
Live imaging reveals impaired neurite growth. **A, B, C,** Individual images derived from live imaging of neurite outgrowth from brain organoid slices indicating the tracks used to quantify neurite behavior. Different conditions including the respective WTs are shown for SOX11 HET (A), MID1 KO (B), and TCF4 HET (C). Scale bars = 200 µm. **D,** Average growth speed depending on the time increment referred to as instantaneous speed as shown by line plots. **E,** End-to-end distances shown by line plots. The dashed lines are fits of the form *a* + *b*√𝑡 (see main text for details). **F,** Line plot indicating the tortuosity, being the ratio of the path length *L* to the end-to-end distance T=L/d, with lower values for T indicating more directed growth. **G,** Mean squared displacement (MSD). **H,** Velocity autocorrelation function (VACF) as a function of the lag time (inset shows the same in log-linear scale and the solid lines show the double exponential fit (see text for details). For **D-H**, Left panel shows SOX11, middle panel MID1, and right panel TCF4 conditions. Error bars indicate standard error of mean. Track numbers: SOX11 WT, 309; SOX11 HET, 233; MID1 WT, 211; MID1 KO, 72; TCF4 WT, 239; TCF4 HET, 199. **I,** Persistence times given in minutes for different conditions. The values were obtained as the larger of the two values from the double exponential fit to the velocity auto-correlation function (shown as solid lines in the insets in panels **H**).

### Molecular profiling of brain organoids unravels molecular nexuses

We next used single cell RNA sequencing (scRNA-seq) to unravel alterations in the molecular framework of neurons associated with the observed cellular defects. We profiled both day 30 and day 60 organoids and embedded the transcriptomes of single cells together (Figures 3A, B, Figures S3A-C). As expected haploinsufficiency of the TFs SOX11 and TCF4 resulted in the downregulation of their respective regulons (Figure S3D). Interestingly, in line with previous results^18^, lower levels of SOX11 and TCF4 also impacted on the regulon activity of the respective other TF. Thus, we found decreased SOX11 regulon activity in TCF4 HET samples (progenitors and neurons) and a lower TCF4 regulon activity in the SOX11 HET samples. In the MID1 KO cells TCF4 regulon activity was found to be increased (Figure S3D). Differentially expressed (DE) genes in neurons across conditions were examined (Figure 3C). In line with the function of MID1 as ubiquitin ligase, we detected an increased expression of genes in MID1 KO neurons (Figures S3E, F) and progenitors (Figure S3G) compared to the respective controls in the MID1 WT organoids. In contrast assessment of DE genes in neurons (Figures S3E, F) and progenitors (Figure S3G) caused by the absence of the TFs SOX11 or TCF4, resulted in more downregulated genes. However, in neurons where we detected a similar cellular phenotype, only 17 genes were commonly deregulated across conditions. Of those only 6 showed deregulation in the same direction (Figure 3D). Interestingly, 4 of these downregulated genes (*MEIS2, HIST1HCH4 (H4C3*^39^*, RPL13A, DPP6*) were previously described as being NDD-associated. Moreover, for *DPP6* a role in the formation of neuronal extensions was shown before^40^. When we moved beyond the individual gene level, to assess deregulated gene regulatory networks, we found an overlap of 20 shared regulons in neurons, the vast majority of which showed the same behavior (Figures 3E, F), i.e. down- or upregulated across experimental conditions. Based on our observations made through live imaging of neurite outgrowth, namely decreased speed and increased tortuosity across conditions, we wondered whether on the level of gene ontology (GO) terms we would find indications on shared deregulated biological processes. We therefore computed the GO terms associated with the DE genes of each individual condition, i.e. in the MID1 KO, the TCF4 HET and the SOX11 HET condition and here found 5 common GO terms (Figure 3G), providing strong evidence for decreased mitochondrial metabolism of the neurons. Further supporting this notion, we detected a considerable convergence on metabolic pathways implicated in the interaction of neurites with the extracellular environment such as arachidonic acid metabolism, eicosanoid metabolism, glutathione metabolism, leukotriene metabolism, and omega-3 fatty acids metabolism (Figures 3H, S4A).

**Figure 3.**
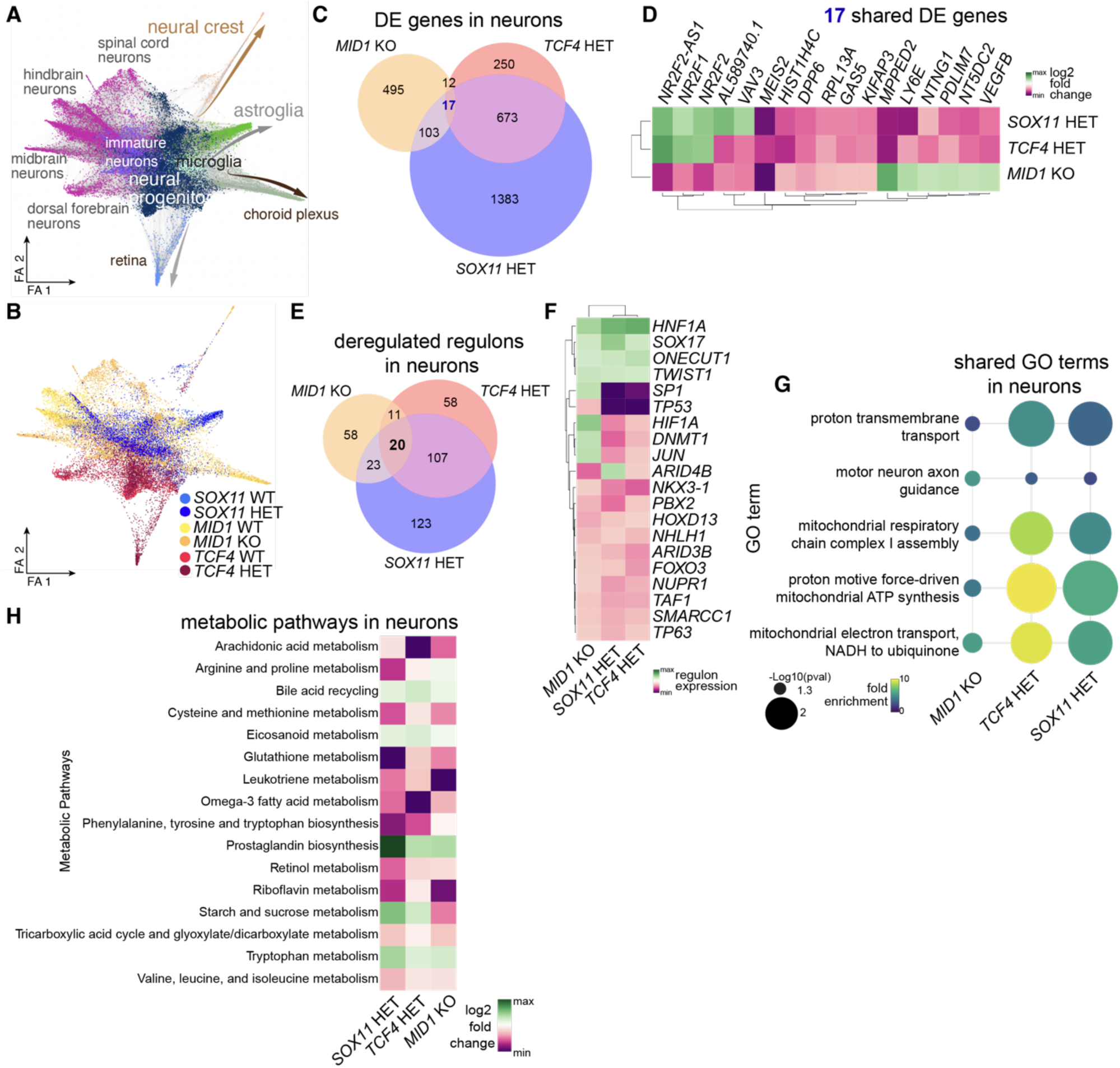
Molecular profiling of brain organoids unravels molecular nexuses. **A,** Force-directed graph embedding of transcriptomes of cells derived from d30 and d60 brain organoids. Distinct lineage trajectories are indicated on the plot. Cells total: 25031; SOX11 WT: 3117; SOX11 HET: 5166; TCF4 WT 3527; TCF4 HET: 3655; MID1 WT: 4191; MID1 KO: 5375. **B** Embedding showing the distribution of cells from different hPSC lines. **A, B,** FA refers to force atlas. **C,** Euler diagrams showing the number of deregulated DE genes (both up and down) in neuros. **D,** Heatmap showing the degree of deregulation (log2 fold change) of the 17 shared deregulated DE genes compared to the respective WT samples across conditions. **E,** Euler diagrams showing deregulated regulons in neuros across the experimental conditions and the respective overlaps. Note the common deregulation of 20 regulons. **F,** Heatmap highlighting the degree of expression of the 20 shared regulons (as determined in E) across conditions. The dendrogram indicates similarities. **G,** Dotplot showing the fold enrichment (indicated by color) of the 5 shared GO terms associated with the DE genes. Significance is indicated by the size of the dot. **J,** Heatmap showing the extent (log2 fold change) of deregulation of metabolic pathways across conditions.

### Assessment of cellular communication uncovers impaired interactions with ECM

Next, we aimed to leverage the scRNA-seq data for a better understanding of putatively impaired cell communication across experimental conditions and therefore determined cell communication probability between cells (signaling from neurons and progenitors to neurons) across conditions using CellChat^41^. Based on the differential expression of signaling pathway members, i.e. ligands and receptors in both progenitors and neurons in each condition, we then computed the upregulated pathways in the respective HET or KO condition compared to their WT controls, discriminating from which cells the signaling originates (Figure 4A). The same analysis was performed for downregulated pathways for which neurons were selected as targets (Figure 4B). Overall, we detected *BMP* as well as 7 pathways implicated in the interaction of cells with the environment/extracellular matrix (ECM) (*EPHA, EPHB, SEMA6, ADGRL, CNTN, NRXN, PTPR*) as shared across the conditions (Figure 4C). Comparing the percentage of differential components shared at the different levels highlights that convergence is stronger at more integrative cellular levels increasing from genes to regulons to cellular communication (Figure 4D).

**Figure 4.**
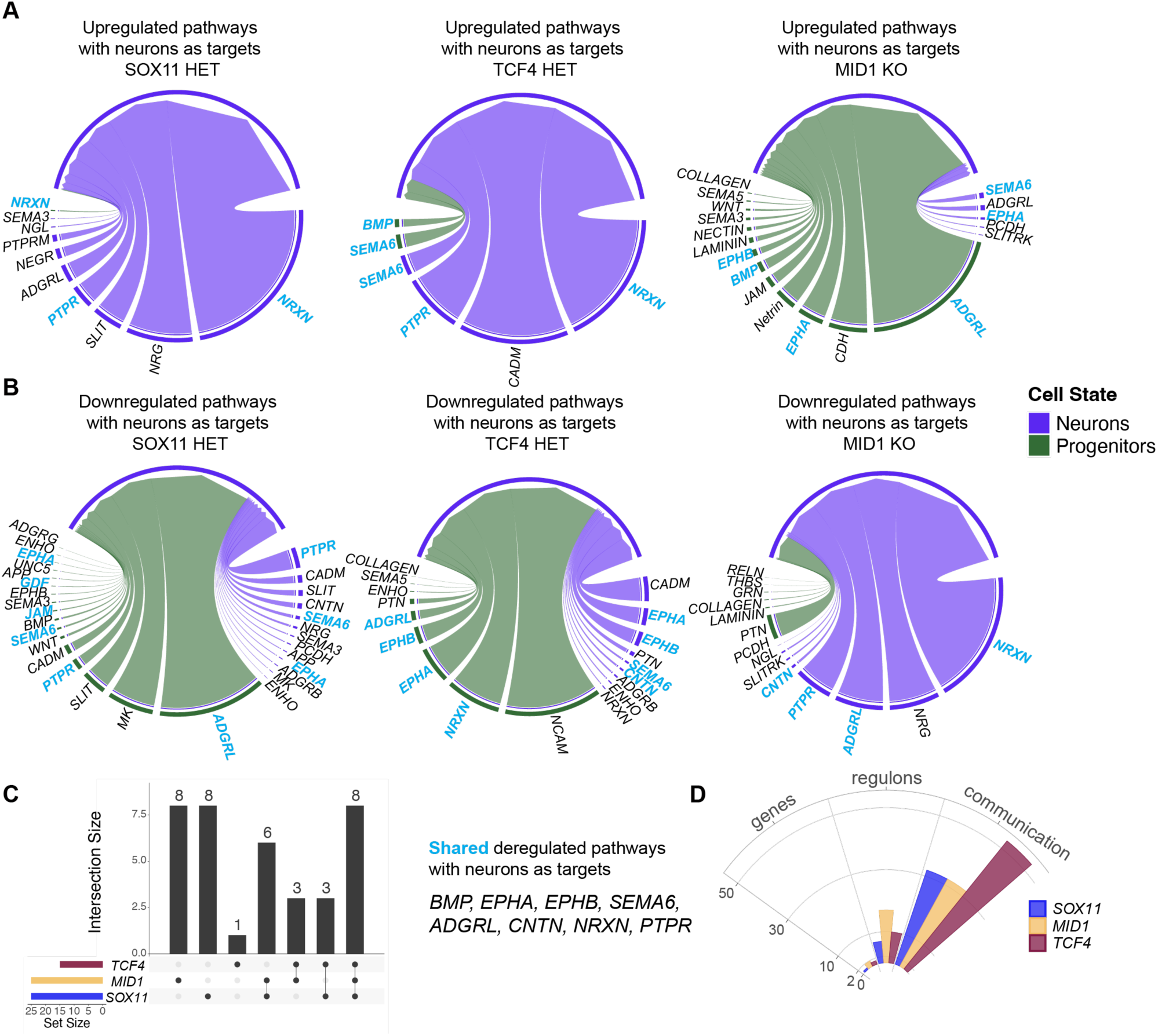
Assessment of cellular communication uncovers impaired interactions with ECM. **A,** Chord diagrams showing upregulated pathways with neurons as targets. Signaling from outside to inside indicating upregulated signaling comparisons. Left panel = SOX11 WT vs SOX11 HET, middle panel = TCF4 WT vs TCF4 HET, and right panel = MID1 WT vs MID1 KO. **B,** Chord diagrams showing downregulated pathways with neurons as targets. Signaling from outside to inside indicating upregulated signaling comparisons. Left panel = SOX11 WT vs SOX11 HET, middle panel = TCF4 WT vs TCF4 HET, and right panel = MID1 WT vs MID1 KO. **C,** Upset plot indicating the number of deregulated signaling pathways including the overlap shared between conditions. The 8 deregulated signaling pathways are listed. **D,** Polar bar graph showing the percentage of the number of shared genes, regulons, and signaling pathways across conditions.

Based on the surprising correlation of phenotypic convergence on a cellular level describing neurite dynamics and molecular convergence highlighting altered interactions of neurites with their environment we next wondered whether changes in neurite-ECM interaction are sufficient to recapitulate the mutant phenotypes. To this end, we rationalized our observation on the statistics of the growing neurites, and put forward a simple phenomenological, two-dimensional *in-silico* model of neurite growth incorporating parameters of neurite-ECM interaction.

### Numerical simulations of neurite growth allow predictions on aberrant cellular parameters

While to date highly complex, multi-scale models of neurite growth exist^42^, our goal was to provide a minimal mechanistic model that focuses on neurite-ECM interaction and provides growth phenotypes as described by our statistical analysis of the tracking data. In our numerical model (Figures 5A, B) neurites are protruding out of the organoid (brown) and are represented by a chain of beads (yellow) connected by harmonic springs. Neurites grow in the extracellular space occupied by particles (grey) representing the extracellular matrix (ECM). The leading bead of the neurite (blue) can establish a temporary link with the matrix particles (magenta), which is randomly chosen within a certain cone angle 𝜃_*link*_ in front of the neurite and in a certain range 𝑅_*link*_. The linked matrix particle then connects to the leading bead by a harmonic spring pulling the leading bead and the ECM particle towards each other. As the neurites advance through the matrix, the geometry of the neurite and positions of the individual matrix particles are altered due to excluded volume interactions when beads overlap (supplementary movie M7 and M8). Neurite growth is composed of two distinct growing modes, i) linear growth independent of ECM interactions and ii) ECM-dependent traction force growth which is the major contributor to neurite extension in our model. Details of the conceptual logic and the equations governing the numerical model are provided in the Methods section.

**Figure 5.**
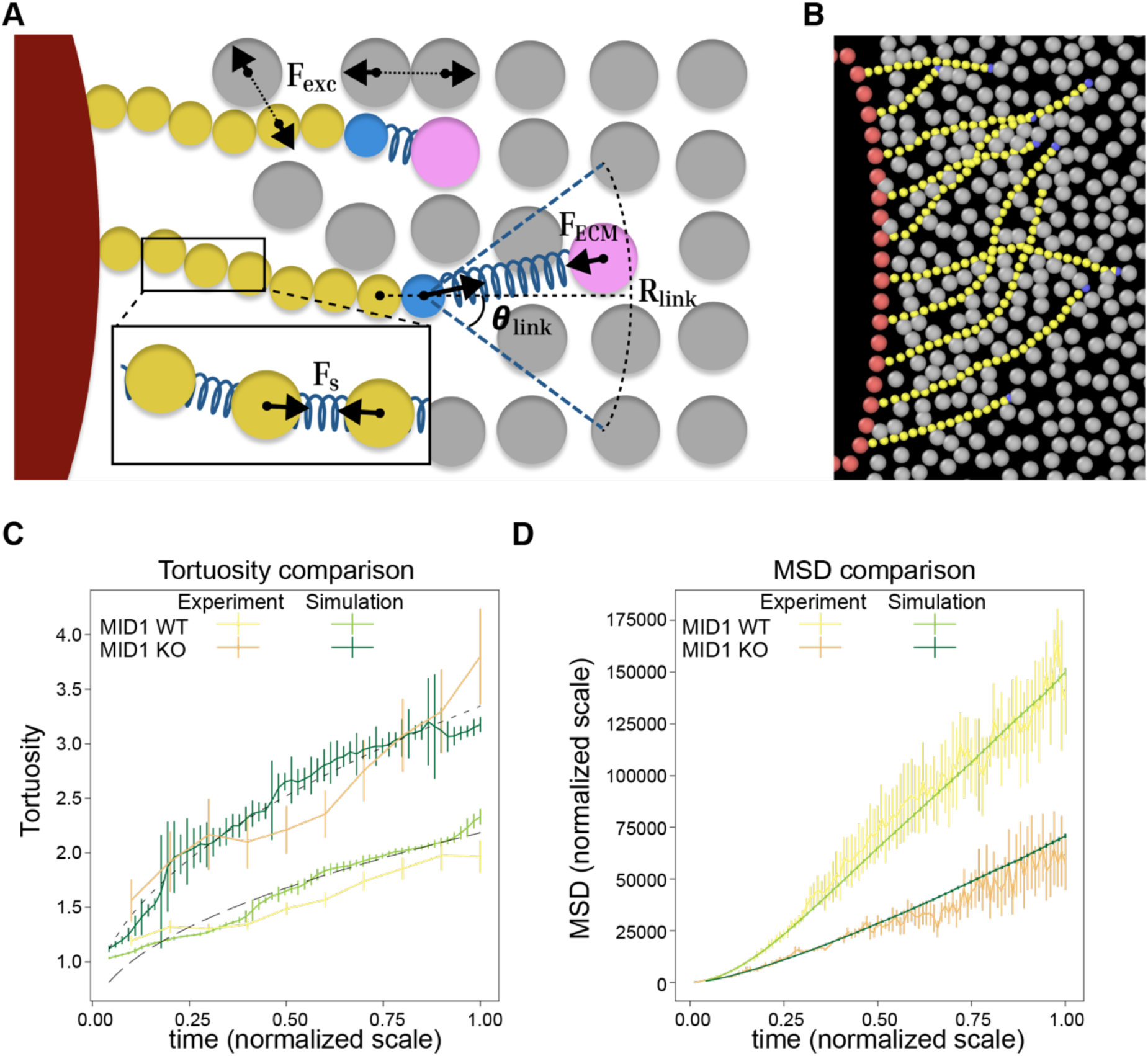
Numerical simulations of neurite growth allow predictions on aberrant cellular parameters. **A,** Schematics of the numerical model. The neurites are represented by a chain of beads (yellow) connected by springs (see the zoom in panel) protrude out of the organoid (brown) and grow in the extracellular space occupied by particles (grey) representing the extracellular matrix (ECM). The leading bead of the neurite (blue) can establish a temporary link with the matrix particles (magenta), which is randomly chosen within a certain cone angle 𝜃_*link*_ in front of the neurites and in a certain range 𝑅_*link*_. The linked matrix particle then connects to the leading bead by a harmonic spring. Meanwhile, as the neurites advance through the matrix its geometry is slowly deformed by repulsive excluded volume interactions. **B,** Snapshot of the actual numerical output of the model. Red beads indicate the boundary of the organoid, yellow beads are the beads of the neurite with a blue bead at its leading end. Grey particles represent the ECM. **C,** Line plots indicating the tortuosity of the neurites. We reproduce higher tortuosity for the MID1 KO as compared to MID1 WT. In the simulation we used the values of 𝜃_*link*_ = 75° for MID1 KO and 𝜃_*link*_ = 30° for MID1 WT. The dashed/dotted lines represent the fit of the simulation data of the form *a* + *b*√𝑡 motivated by the theory of random walks (see the text for details). Specifically, it is 0.27 + 3.34√𝑡 and 0.61 + 1.48√𝑡 for MID1 KO and WT respectively. **D,** MSD as a function of time as shown by line plots. Simulations reliably recapitulate the experimental data. For **C, D,** Error bars indicate standard error of mean.

To apply our model to the experimental data we decided to focus on the MID1 condition as it shows most pronounced differences between WT and KO phenotypes. We first searched for parameters that generate neurite growth trajectories with matching values of tortuosity for the MID1 WT (see Table S1). To compare the numerical simulations and real data we rescaled both the time axis of the experiment and simulations to the value of 1 (Figure 5C). We also generated the respective plots for the MSD and normalized the *y* axis of the plot to match the experimental data (Figure 5D). Our simulations are scale-agnostic, and by rescaling the MSD values we fixed the spatial scale in the simulations. With these steps we now set the time and space scales of the WT data. Next, we modulated the parameters of the simulations, but not their time and space scaling, to match the mutant data. Remarkably, changing a single parameter, i.e. the opening angle in which the neurite can choose an ECM particle for the traction force realization, is sufficient to transform the MID1 WT to the MID1 KO data both for the MSD and tortuosity (Figure 5D). Here we also note that in agreement with the random walk theory, the tortuosity being the ratio of track length (growing proportional to time) and end-to-end distance (scaling as square root of time) grows as a square root of time. Both, our simulations and experimental data follow this trend as indicated by dashed lines in Figure 5C.

Our numerical simulations of neurite growth have shown that it is not only possible to interpret the growth of neurites following the theory of random walks but moreover allow to predict the underlying parameters sufficient to recapitulate the diseases phenotypes of neurite growth. In sum, we show that three distinct neurodevelopmental diseases sharing aberrations in long-range connections as a common phenotype converge on the interaction of the growing neurite with its environment.

## Discussion

Employing human PSCs to generate brain organoids has enabled a tremendous leap forward to model human NDDs^43^. Here we utilized brain organoids to tackle the question whether distinct NDDs sharing aberrations in long-range connections as a common phenotype converge on molecular and cellular nexuses, responsible for the establishment of neuronal projections. In a comparative assessment we have generated brain organoids from hPSCs with mutations in either the gene *SOX11* causing CSS9, the gene *MID1* causing OS, and the gene *TCF4* causing PTHS, and their respective isogenic controls. Tissue phenotyping revealed a transient delay in neuron formation in the SOX11 HET and the MID1 KO organoids, likely caused by a change in their progenitor population (Figures 1, S2). While SOX11 HET progenitors showed lateral expansion, the MID1 KO progenitors were found to expand radially, which is in line with previous work^16,35,44^. Importantly, the proliferation phenotypes were transient in both lines resolving at later stages with all lines showing no difference in the balance between proliferative SOX2+ areas and differentiating MAP2+ areas at later stages. TCF4 HET organoids showed a neuronal migration phenotype with a transient accumulation of DCX positive cells in the VZLS reminiscent of periventricular heterotopias and in line with previous studies highlighting the role of TCF4 in migration^45,46^. However, these DCX+ nodules in the VZLS resolved over the course of organoid development. Thus, neither the balance between proliferation and differentiation of neural stem cells nor the migration of young neurons are shared cellular phenotypes between the three NDDs. Concomitantly, also on a molecular level the genes and gene regulatory networks that are affected in neural stem cells by the individual mutations show minimal overlap. Hence, our data suggest that in neural stem cells the cellular and molecular consequences of the mutations are specific to each NDD and show little to no convergence (Figures 1, 3, S2, S3).

We next turned our attention to the neurons produced by the neural stem cells and in particular to neurite growth^47–52^ essential for the proper formation of neuronal long-range projections. Using live-cell imaging to follow neurite growth over the course of three days we functionally assessed the impact of the three NDD causing mutations on neurite growth dynamics. We established that the theory of persistent random walks^53^ is a useful framework to quantify neurite growth behavior. Employing such quantifiers revealed that neurite growth dynamics showed a surprising consistency in phenotypes across conditions. These analyses unraveled that the SOX11 HET, MID1 KO, and TCF4 HET neurites grew slower and more awry, i.e. with an increased tortuosity. All neurites, wildtype and mutants, showed a remarkable persistence in their growth direction early in their trajectory, i.e. the growth direction correlated with the previously traveled growth direction of the neurite. With time of growth, and distance from the soma, this correlation decays, eventually resulting in almost random directionality choices. Strikingly, all mutations resulted in neurites which lost the correlation in directionality much earlier than their controls, i.e. they showed a decreased persistence time. Our data hence provides a cellular explanation why in all three NDDs long-but not short-range projections are affected (Figure 2).

Finding this striking convergence of phenotypes on a cellular level we wondered whether there are molecular correlates in neurons showing similar convergence. To dissect the underlying molecular framework for the neurite phenotypes, we sampled brain organoids of all conditions for scRNAseq and analyzed their neurons. While we found very few shared deregulated genes across the affected neurons (SOX11 HET, MID1 KO, TCF4 HET), we observed a gradual increase in the degree of convergence from deregulated genes to deregulated gene regulatory networks to deregulated cellular communication. This suggests that the mutations in the three different NDD causing genes initially trigger very different molecular responses that through the organizational principles within the cell converge on similar functional deficits at more integrative levels. When we then interrogated which cellular communication systems are specifically affected, we found that in particular the interaction of neurons with their extracellular environment is altered in mutant neurons (Figure 4). Interestingly, when scrutinizing the difference in cellular metabolism of neurons induced by the mutations, we found changes in arachidonic acid metabolism, eicosanoid metabolism, glutathione metabolism, leukotriene metabolism, and omega-3 fatty acids metabolism, all linked to ECM and neurite growth^54–65^.

To test whether alterations in the neuron-extracellular environment interaction can explain the mutant phenotypes we developed a phenomenological, two-dimensional model of neurite growth incorporating the molecular interaction of neurites with the extracellular matrix (ECM). Using this numerical model and iterating the different parameters revealed that the modulation of a single factor was sufficient to recapitulate the mutant phenotype from wildtype data. The *in-silico* model suggests that the angle (𝜃) under which the growing neurite chooses its next ECM interaction point is crucial not only to recapitulate the mutant tortuosity graphs but also the MSD data describing the exploratory behavior of neurites which is essential for the formation of long-range projections. Employing the theory of random walks^36^ the opening angle 𝜃 can be related to the persistence time 𝜏 via 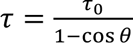, where 𝜏_0_ can be interpreted as the time required for the growth of a new neurite segment. This explains how different tortuosity in the mutant data can be linked to changes in the angle 𝜃 under which a neurite interacts with the ECM and explains how altered ECM-neurite interaction is responsible for changes in the persistence of the trajectories. Given that short-range connecting neurons require a considerably shorter persistence time to reach their target, our *in-silico* model of neurite growth provides a conceptual basis explaining why such changes in the neurite-ECM interaction affect predominantly long-but not short-range neuronal connections.

To conclude, we show that three NDDs CSS9, OS, and PTHS caused by mutations in *SOX11*, *MID1*, and *TCF4* respectively sharing aberrations in long-range connection as a common phenotype converge on the interaction of the growing neurite with its environment. Strikingly, our data indicate that in neurons across all conditions metabolic pathways implicated in the interaction of cells with their environment are altered. Thus, our findings open up the intriguing possibility that extrinsic modulation of the cellular metabolism can modulate the severity of disease phenotypes.

## Online Methods

### Human pluripotent stem cells used throughout the study

Pluripotent stem cell lines were used throughout the study. In detail, we used the human embryonic stem (hES) cell lines H9 to produce the SOX11 lines and the hES cell line Hues6 to generate the TCF4 lines. The human induced pluripotent stem cell (hiPSC) line S-21 derived from a healthy male individual was used to generate the MID1 lines. The generation of the S-21 hiPSC line from male fibroblasts was described earlier^35^. Fibroblasts were acquired at the University Medical Center in Mainz following approval by the local ethical committee (No. 4485). Consent for further analysis and usage for research in an anonymized way was given.

### Genome editing

Generation of genome edited hES lines was done as described earlier ^16^. To generate the SOX11 heterozygous (HET) cell lines, the human embryonic stem cell (hES) line H9 was used. Targeted editing of the SOX11 gene, located on chromosome 2, focused on Exon 1, using the following oligonucleotide sequences to clone the gRNA forward 5’-CACCGGacgccgtcttgcaccagtc-3’ and a reverse 5’-AAACgactggtgcaagacggcgtcC-3’. The SOX11 WT line corresponds to clone H9-57, while the clone H9-16 is the SOX11 HET line carrying a 7bp insertion.

The TCF4 HET line was derived from the hES line Hues6. To target Exon 18 of the TCF4 gene located on chromosome 18q21.2, we used the following oligonucleotide sequences to clone the gRNA forward 5’-CACCGGgggcttgtcactcttgagg-3’ and the reverse 5’- AAACcctcaagagtgacaagccccC-3’. The TCF4 WT line is designated as clone H6-7, and the TCF4 HET line is clone H6-2 carrying a 11bp deletion.

Validation of CRISPR editing in SOX11 and TCF4 was conducted using ICE analysis. For SOX11, ICE analysis demonstrated a high model fit (R² = 0.92), with primary indel contributions indicating a 7-bp insertion in 45% of alleles. For TCF4, ICE analysis revealed a high model fit (R² = 0.87) with 51% contribution of a 11bp deletion.

### Quantification of protein levels in hPSCs

The quantification of TCF4 protein levels was done as described earlier^28^. In short, multichannel images for DAPI and TCF4 were acquired from 2D hPSCs cultures using an EVOS M7000 Imaging System (Thermo Fisher). The DAPI channel was used to segment the nuclei using celldetection (v.0.4.9)^66^ with the pretrained ginoro_CpnResNeXt101UNet-fbe875f1a3e5ce2c model. In each segmented nuclei we then quantified the TCF4 protein levels employing measure.regionprops of the skimage package (v0.25.2) in Python (v3.12.9). The mean intensity was plotted.

### Brain organoid formation and processing

mTeSR™ Plus medium (StemCell Technologies) was used to culture all hPSC lines used in this study. Cells were grown on Matrigel-coated dishes in 5% CO_2_ at 37°C until a confluency of 80-90% was reached. Brain organoid formation was used according to a published protocol including small adaptations ^67^. Briefly, accutase (Thermo Fisher) was used to generate single cell suspensions of hPSCs. Following centrifugation, cells were resuspended in organoid formation medium supplied with 4ng/ml of low bFGF (Peprotech) and 5 µM ROCK-inhibitor Y-27632 (StemCell Technologies). Organoid formation medium consisted of DMEM/F12 + GlutaMAX-I (Thermo Fisher), 20% KOSR (Thermo Fisher), 3% FBS (Thermo Fisher), 0.1 mM MEM-NEAA (Thermo Fisher), 0.1 mM 2-mercaptoethanol (Sigma-Aldrich). 9,000 cells in 150 µL organoid formation medium/well were aggregated in low attachment 96-well plates (Corning) for at least 48 hours during which embryoid bodies (EBs) formed. After 72 hours half of the medium was replaced with 150µl of new organoid formation medium without bFGF and ROCK-inhibitor. At day 5 neural induction medium consisting of DMEM/F12 + GlutaMAX-I (Gibco), 1% N2 supplement (Gibco), 0.1mM MEM-NEAA (Gibco), and 1µg/ml Heparin (Sigma-Aldrich) was added to the EBs in the 96-well plate to promote their growth and neural differentiation. Neural induction medium was changed every two days until day 12/13, when aggregates were transferred to undiluted Matrigel (Corning) droplets. The embedded organoids were transferred to a petri dish (Greiner Bio-One) containing organoid differentiation medium without vitamin A. Three or four days later the medium was exchanged with organoid differentiation medium with vitamin A and the plates were transferred to an orbital shaker (IKA Rocker 3D digital) set to 30 rpm inside the incubator. Medium was changed twice per week. Organoid differentiation medium consisted of a 1:1 mix of DMEM/F12 + GlutaMAX-I (Thermo Fisher) and Neurobasal medium (Thermo Fisher), 0,5% N2 supplement (Thermo Fisher), 0.1 mM MEM-NEAA (Thermo Fisher), 100 U/ml penicillin and 100 µg/ml streptomycin (Thermo Fisher), 1% B27 +/-vitamin A supplement (Thermo Fisher), 0.025% insulin (Sigma-Aldrich), 0.035% 2-mercaptoethanol (Sigma-Aldrich). For fixation, organoids were transferred from petri dishes to 1.5 ml tubes. Organoids were washed with PBS and then fixed with 1xPBS buffered 4% paraformaldehyde (PFA, Carl Roth) for 30 min. Time of PFA fixation was extended up to one hour depending on the size of the organoids. Afterwards, organoids were washed three times for 10 min with PBS and incubated in 30% sucrose (Sigma-Aldrich) in PBS for cryoprotection. For cryosectioning, organoids were embedded in Neg-50™ Frozen Section Medium (Thermo Fisher) on dry ice. Frozen organoids were cryosectioned in 30 μm sections using the Thermo Fisher Cryostar NX70 cryostat. Sections were placed on SuperFrost Plus™ Object Slides (Thermo Fisher) and stored at -20°C until use. To control for batch-to-batch variation we started the generation of organoids of different conditions (SOX11 HET, TCF4 HET, and MID1 KO) together with their controls and considered them as one batch. The analysis of a given phenotype is always batch-controlled, i.e. normalized to the mean value of the controls in the respective batch thereby minimizing the impact of batch-to-batch heterogeneity and focusing on the consequences of the mutations.

### Immunocytochemistry

Frozen organoid slices were thawed and washed once with PBS. Post-fixation of organoid slices was achieved using 4% PFA for 15 min followed by three washing steps with PBS for 5 min. During the entire staining procedure, slides were kept in humidified staining chambers in the dark. For antigen retrieval, the sections were incubated for 20 minutes at 70°C in HistoVT One buffer (Fisher Scientific), diluted 1:10 in distilled water. Slices were then washed briefly with blocking solution (PBS, 4% normal donkey serum (NDS, Sigma-Aldrich), 0.25% Triton-X 100 (Sigma-Aldrich)) followed by 1 h incubation with blocking solution at room temperature. Primary antibodies were diluted in antibody solution (PBS, 4% NDS, 0.1% Triton-X 100) and tissue sections were incubated overnight at 4°C. Next, following three washes using PBS with 0.5% Triton-X 100, secondary antibodies were added diluted in antibody solution and incubated for 2 h at room temperature. Sections were washed three times with PBS containing 0.5% Triton-X 100 for 5 min. Slides were counterstained with DAPI 1:1000 in PBS for 5 min followed by one washing step with PBS. Lastly, organoid sections were mounted using Aqua Polymount (Polysciences).

Antibodies used were selected according to the antibody validation reported by the distributing companies. The following primary antibodies were used: rabbit anti-ARL13B (Proteintech, 17711-1-AP, IgG, 1:250), mouse (IgG1) anti-MAP2 (Sigma-Aldrich; M4403; IgG1, 1:300), rabbit anti-SOX2 (Abcam; ab137385; IgG, 1:300), human anti-PAX6 (Miltenyi Biotec, 130-107-582, 1:300), mouse anti-N-cadherin (BD Bioscience, 610921, IgG1, 1:300), sheep anti-TTR (Biorad, AHP1837, IgG, 1:100), mouse anti-PH3 (Cell Signaling Technologies, 9706, IgG1, 1:300), guinea pig anti-DCX (Merck Millipore, AB2253, 1:300), mouse anti-OCT4 (Santa Cruz, sc-5279, IgG2a, 1:300), goat anti-Nanog (R&D Systems, AF1997). The following secondary antibodies were used: goat anti-rabbit Alexa 488 (Thermo Fisher, A11008, 1:500), donkey anti-sheep Alexa 488 (Thermo Fisher, A11015, 1:1000), goat anti-mouse IgG1 Alexa 555 (Thermo Fisher, A21127, 1:500), goat anti-rabbit Alexa 633 (Thermo Fisher, A21070, 1:500), goat anti-chicken Alexa 488 (Thermo Fisher, A11039, 1:1000), goat anti-mouse IgG1 Alexa 647 (Thermo Fisher, A21240, 1:1000), goat anti-rabbit Alexa 647 (Thermo Fisher, A21245, 1:1000), anti-rat Cy3 (Thermo Fisher, A10522, 1:1000), goat anti-guinea pig IgG Alexa Fluor 555 (Thermo Fisher, A21435, 1:1000).

### Microscopy and image analysis

Epifluorescence pictures were taken using the EVOS^TM^ M7000 Imaging System (Thermo Fisher). Z-resolved pictures were acquired using an Apotome 2 (Zeiss) equipped the Colibri5 light source (Zeiss). To quantify the VZLS area in each organoid section, we measured both the total organoid area (excluding cyst-covered regions) and the VZLS-covered area using FIJI (v1.52-1.53) selection tools. The fraction of organoid area occupied by VZLS was then computed in R (v3.5.1-4.1.2), averaging values across different sections from the same organoids and normalizing them to the mean value of control organoids within each batch.

For quantification of SOX2- and MAP2-covered areas in organoid sections, epifluorescence images obtained with the EVOS M7000 were processed using OpenCV (v4.4.0 - 4.5.1) in Python (v3.9.1-3.9.10) for automated thresholding and pixel counting. The neural area was determined based on the number of thresholded pixels corresponding to SOX2 or MAP2, followed by calculating the SOX2/neural area and MAP2/neural area fractions using NumPy (v1.21.5) and pandas (v1.3.4). For segmentation of pH3-positive cells, the celldetection package (v0.4.9) was used.

### Live imaging of neurite outgrowth

For live imaging, organoids at day 60 were dissected by hand into fine slices using a sterile scalpel. The slices were then retrieved with a plastic Pasteur pipette and placed on Matrigel-coated 6-well plates with a minimal amount of medium, allowing them to adhere for 5-6 hours before adding additional medium. Five hours prior to imaging, SiR-tubulin, a fluorogenic microtubule labeling probe (Tebubio, SC002), was added to the slice culture at a final concentration of 100 nM to label neurites and enable visualization of neurite outgrowths. The plate containing the organoid slices was then placed in the EVOS™ M7000 Imaging System (Thermo Fisher), equipped with a humidity chamber to maintain 37°C, 70% humidity, and 5% CO_2_. Live imaging was conducted over a 72-hour period with images taken every 20 minutes.

### Neurite tracking

Individual live imaging pictures were assembled into movies using a custom FIJI (v1.52-1.53) macro developed in-house. Individual neurites were tracked manually using the MtrackJ (v.1.5.2) plugin in ImageJ, to follow the growth of each neurite tip frame by frame.

### Statistical analysis of the neurite growth

Neurite tracking provided us with a data set describing the two-dimensional coordinates of the growing neurite tip at discrete timepoints points *i* for every trajectory *j*: {*t_i_,x_j,i_, y_j,i_*}. The pixel coordinates are then converted to micrometers (1 pixel = 0.5029μm). We first used these coordinates to plot and inspect the traced trajectories. We discarded trajectories that showed sudden big jumps in displacement (>110μm between two timeframes). If the tip was not moving for more than 200 min, the track was terminated at that point. Tracks starting at later times relative to the start of the movie were shifted to the starting point at t=0. In total we obtained the following numbers of trajectories after filtering: SOX11 WT (309), SOX11 HET (233), MID1 WT (210), MID1 KO (73), TCF4 WT (240), and TCF4 HET (199). All the trajectories used for the final analysis can be accessed via Zenodo 10.5281/zenodo.15213837.

We first quantified the instantaneous growth speed of neurites, which is defined as:

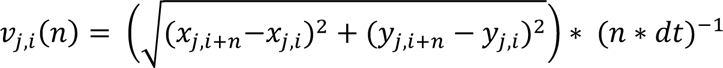

where *dt=*20 min is the temporal resolution of our data, and *n* denotes how many time steps are used to calculate the instantaneous speed; we used the values of *n* in the range from 1 to 10 (Figure 2D). We averaged these values along the trajectory and for all trajectories of the same condition. We next evaluated the length of the path traveled by the tip as a function of time (*t*=*N*dt* after *N* steps):

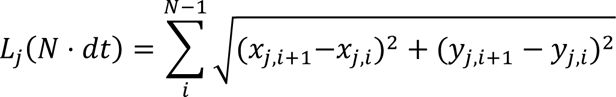

and averaged that over all trajectories *j* of the condition. Note that trajectories are of different lengths, so generally for larger times there are fewer trajectories contributing to the average. End-to-end distance was calculated using:

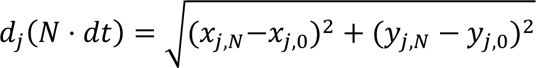

With the general definition of the Mean Squared Displacement 𝑀𝑆𝐷(𝑡) =< (𝒓(𝒕) − 𝒓(𝟎))^𝟐^ > for a discrete set of trajectories this results in:

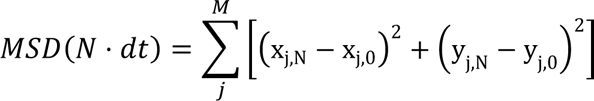

The velocity autocorrelation function is defined as VACF(t) = < v(t_0_ + t) ⋅ v(t_0_) >. We used the shortest time interval *dt* to define the vectors of velocity and look at the average scalar product of velocities along the same trajectory but shifted by the time *t*. Assuming that the tip growth is a steady process, we can additionally perform averaging over the starting point *t*_0_ along the trajectory (time averaging). As a result, we obtain a decaying function with two distinct regimes (see Figure 3H): rapid decay for very short times (of the order of our temporal resolution) and a slower decay. The first regime shows the loss of correlation due to rapid velocity fluctuations, while the slower decay corresponds to the persistence of growth. We use a simple fitting function Ae^−𝑡/τ_1_^ + Be^−t/τ_2_^ and focus our attention on the slowest time scale 𝜏_2_ that we call a persistence time. These times are provided in Figure 3I.

### Single cell RNA-seq data generation

For the scRNA-seq experiment, organoids were dissociated using the Neural Tissue Dissociation Kit P (Miltenyi Biotec). Briefly, selected organoids were cut into smaller pieces, washed with medium, and three times for 5 minutes with 1xPBS. Organoid pieces were transferred to a tube containing the enzyme mix P (according to the manufacturer’s protocol) and incubated at 37°C for 10 minutes. Pieces were then triturated gently with a 1000p pipette tip and incubated for another 10 minutes at 37°C in the presence of enzyme mix A (according to the manufacturer’s protocol). Pieces were then triturated gently with a 1000p and a 200p pipette tips and incubated for 5 minutes at 37°C. Cell suspension was filtered with a 30 µm filter (Miltenyi Biotec) and centrifuged at 300xg for 5 minutes. After a second filtration step with a 20µm filter (Miltenyi Biotec) and subsequent centrifugation as described above cell pellet was resuspended in 100µl 1xPBS (without Ca^2+^ and Mg^2+^). Cells were counted and tested for viability with Trypan Blue and the automated cell counter Countess (Thermo Fisher). Cells were diluted to an appropriate concentration to obtain approximately 5000 cells per lane on a 10X Next GEM chip G v3. Libraries were constructed according to the Chromium Single Cell 3’Reagent Kit User Guide (v3 Chemistry Dual Index) of 10X genomics and sequenced on an Illumina NovaSeq 6000.

### Single cell RNA-seq data preprocessing, clustering, visualization

De-multiplexing, genome alignment to the GRCh38 reference genome, and sequencing depth normalization were performed using the “count” and “aggr” functions of Cell Ranger (10x Genomics, versions 4.0.0 to 6.0.2). Further preprocessing, clustering, embedding, and visualization were carried out using Scanpy (v1.10.2)^68^. Additional de-multiplexing was performed using the Souporcell package (singularity 4.1.2) ^69^, aligning to the GRCh38 reference genome the four different conditions and also to the H9 hESC sequence using the STAR package (2.7.11b)^70^ as reference for the SOX11 samples, and the Y chromosome-related genes RNA expression as reference for the MID1 samples.

Cells were filtered based on the following criteria: those with fewer than 500 detected genes were excluded, along with condition-specific thresholds for UMI counts, gene numbers, and mitochondrial gene percentages. Specifically, for d30 SOX11 WT and TCF4WT, and d60 MID1 KO cells with UMI counts between 0 and 20,000, gene counts between 1,500 and 6,000, and more than 7.5% mitochondrial genes were removed. In d30 SOX11 HET, TCF4 HET and d60 MID1 WT, cells with fewer than 2000 or more than 20,000 UMI counts, fewer than 1,500 or more than 6,000 genes, and over 7.5% mitochondrial genes were filtered out. For d60 SOX11 WT, TCF4 WT, the filtering thresholds were UMI counts between 2,000 and 20,000, gene counts between 2,000 and 6,000, and over 6% mitochondrial genes. For d60 SOX11 HET and TCF4 HET, cells with fewer than 0 or more than 10,000 UMI counts, fewer than 0 or more than 5,000 genes, and over 6% mitochondrial genes were excluded. For d30 MID1 WT, cells with 7,500 to 25,000 UMI counts, 3,500 to 6,750 genes, and over 7.5% mitochondrial genes were excluded. In the d30 MID1-KO condition, cells with UMI counts between 6,000 and 30,000, 3,000 to 7,500 genes, and over 6% mitochondrial genes were removed. Genes expressed in fewer than five cells were also excluded from the analysis.

Principal component analysis (PCA) was conducted using Scanpy, selecting the 15 nearest neighbors in the top 18 principal components (PCs). Clustering was performed using the Leiden algorithm with a resolution of 1. After clustering, cells from clusters ‘17’, ‘6’, ‘9’, and ‘18’, which corresponded to mesenchymal-related clusters, were excluded. Additionally, X and Y chromosome-related genes were temporarily removed as some cells were clustered based on sex chromosome gene expression rather than cell identity. Once these cells were identified and removed from the unnormalized data, preprocessing was repeated with normalization and log transformation, followed by identification of highly variable genes using default Scanpy settings. Confounding factors such as cell cycle scores, mitochondrial gene percentage, and UMI counts were regressed out. PCA was repeated, followed by selecting the 15 nearest neighbors in the top 16 PCs, Leiden clustering at a resolution of 1.7, and batch integration using bbknn. UMAP embedding was computed with a minimum distance of 0.3 and a spread of 1. Partition-based graph abstraction (PAGA) was calculated with a threshold of 0.22, and force-directed graph embedding utilizing ForceAtlas2 provided by the python package fa2 (v3.5) based on PAGA distances was performed for visualization. Finally, the previously removed X and Y chromosome-related genes were reintroduced for further analysis.

### Single cell RNA-seq, RNA velocity, and pseudotime estimation

To determine spliced and unspliced transcripts, the Cell ranger produced BAM files were sorted by the cell barcode with samtools (v1.10) and counted with velocyto (v0.17)^71^. RNA velocity estimation was then done with scVelo (v0.2.3)^72^. In detail, moments were calculated with the ‘connectivities’ mode on the top 50 PCs and 10 closest neighbors. After recovering the velocity dynamics, latent time was calculated, a measure for the developmental time (pseudotime) exclusively depending on transcriptional dynamics. RNA velocity was then computed, using latent time, differential kinetics and highly variable genes with the stochastic model.

### Differential gene expression and GO term analysis

Differentially expressed genes in progenitor cells (Leiden clusters 0, 3, 4, 5, 7, 10, 16, 21, 22, 24) and neurons (Leiden clusters 1, 9, 12, 13, 14, 17, 18, 23, 26) were identified between SOX11 HET and SOX11 WT, TCF4 HET and TCF4 WT, and MID1 KO and MID1 WT. The analysis was performed using the “rank_genes_group” function of Scanpy, with significance tested by the Wilcoxon rank sum test. Genes with an adjusted P-value < 0.05 and an absolute log2 fold change > 0.5 were considered differentially expressed. Venn diagrams were generated using these differentially expressed genes. For upregulated shared genes, genes with an adjusted P-value < 0.01 and an absolute log2 fold change > +0.5 were included. For downregulated shared genes, those with an absolute log2 fold change < -0.5 were used. Analyses were conducted separately for neurons and progenitor cells. GO analysis was performed using the R (v4.3.2) package TopGO (v2.54.0). All genes expressed in the neuronal or progenitor cell datasets were used as the background. GO terms with fewer than 10 annotated genes were excluded from the analysis. The default “weight01” algorithm was applied to assess GO enrichment. Only GO terms with more than three differentially expressed genes and a P-value < 0.05, as determined by Fisher’s exact test, were considered significant.

### Regulons

We used decoupleR (v1.6.0)^73^ and Omnipath (v1.0.8) python packages to infer transcription factor activity, following decoupler’s tutorial on “Transcription Factor Activity Inference.” The activity inference was calculated using a univariate linear model across the entire dataset with the run_ulm function, applying the default parameters except for the batch_size, which was set to the total sample size (25031). To compare transcription factor activity, we subsetted the dataset for each mutant condition (SOX11 HET, TCF4 HET, and MID1 KO) and used the rank_sources_groups function, where the corresponding WT equivalent served as the reference group. To identify significant shared regulons, we filtered the three resulting datasets for transcription factors with a p-value lower than 0.01 and a mean change greater than 0.2 or smaller than -0.2.

### Metabolic Analysis

We used METAflux (v1.0.0) ^74^ to infer metabolic flux distributions from transcriptomic data. Flux estimation was performed using the tutorial’s default settings, with a bootstrapping procedure of n_bootstrap = 200 to obtain robust flux estimates. To model the mTesR cell medium composition, we custom-generated a cell_medium file using the mTesR media compounds listed in the nutrients_lookup_file. To ensure consistency across experimental conditions, analyses were restricted to Neurons and Progenitors cell types. For statistical evaluation, we computed the cumulative distribution function (CDF) for each reaction and retained only those with a p-value below 0.05.

### Cell-cell communication analysis

To analyze cell-cell communication, we utilized the CellChat R package (version 2.1.2) ^41^. To reduce noise, only neuronal and progenitor clusters were retained, and the data was subset for each condition (SOX11 WT, SOX11 HET, TCF4 WT, TCF4 HET, MID1 WT, and MID1 KO). The CellChat object was generated following the “CellChat-vignette,” with modifications to include all communications from CellChatDB, except for “Non-protein Signaling.” 10% truncated mean as the statistical method was used to infer significant ligand-receptor pairs, allowing us to capture a higher number of interactions while maintaining robust predictions. For the comparison of two CellChat objects, “Comparison analysis of multiple datasets” vignette was followed.

### In-silico model

To rationalize our observations on the statistics of the growing neurites we put forward a simple phenomenological, two-dimensional model of neurite growth where the special attention is put on the neurite interactions with the extracellular matrix (ECM). In particular, we implemented neurite growth which is dependent on the traction forces of the leading edge of the protrusion with the ECM. Below we provide a description of the model and governing equations as well as their numerical implementation.

Each neurite is approximated as a chain of circular beads with radius 𝑅_𝑛_ and mobility 𝜈_𝑛_(Figure 5A). All beads (yellow within chain), except for the foremost bead (blue) along the chain are connected to each other by ideal elastic springs with a spring constant 𝜅_𝑠_ and an equilibrium length 𝑙_0_. Thus, if the distance between the two neighboring beads 𝑙 deviates from 𝑙_0_ a restoring force acts on the beads along the spring direction: 𝐹_𝑠_ = 𝜅_𝑠_(𝑙 − 𝑙_0_). Neurite growth occurs in a space filled by the ECM which was simulated as a collection of beads (grey)^75^ of size 𝑅_𝐸𝐶𝑀_ and mobility 𝜈_𝐸𝐶𝑀_. We chose ECM particles to be larger than neurite particles and with lower mobility. Generally, all beads (neurites and ECM) are modeled as non-overlapping particles (only beads of the neurites are allowed to cross each other to mimic 3D situation). The overlap is prevented by the excluded volume force which only acts when to beads overlap (|*r_i_* − *r_j_*| < (*R_i_* + *R_j_*)) :*F_exc,ij_* = −𝜖 𝛴 (|*r_i_* − *r_j_*| − (*R_i_* + *R_j_*)) x *n_ij_*, where, *r_m_* and *R_m_* are position vector and radius of the *m*-th bead respectively, 𝒏_𝑖j_ is the unit vector connecting the pair of beads *i* and *j*, and 𝜖 is the spring constant chosen to be higher than the rest of the spring constants to replicate hard-body repulsion.

Neurite growth is simulated as an extension of the foremost bead (blue) and captures two distinct mechanisms: i) linear growth independent of ECM and ii) ECM-dependent traction force growth. For linear growth i) neurite extension starts by adding a new bead at the position of the foremost bead. The new bead is connected to the existing neurite by a spring with the same spring constant 𝜅_𝑠_, but its equilibrium length starts to grow linearly with time: 𝑙_g_ = 𝛼𝑡, with the growth rate 𝛼. When 𝑙_g_ reaches 𝑙_0_, the growth stops, the next new bead is added, and the process repeats. This mechanism describes slow, ECM-independent growth of the neurite playing only a minor role in our model. However, the ECM-dependent traction force mechanism (ii) is the major contributor to neurite extension in our model. Here, the new bead is also added at the position of the previous leading bead. The new bead is connected to the existing neurite by a weaker spring with the spring constant 𝜅_𝑡𝑟*a*𝑐_ and a slowly growing equilibrium length describing a slow background neurite growth in the absence of the matrix. The new bead now randomly selects an ECM bead in a cone with an opening angle 𝜃_*link*_ along the direction of the two trailing beads. Furthermore, the distance to the selected ECM bead is limited by the range 𝑅_*link*_. The ECM bead (magenta in Figure 5A) and the leading bead (blue) are connected by the temporary spring with a certain lifetime mimicking the finite lifetime of adhesive interaction pulling the blue bead forward. The force between the leading bead and the ECM is thus given by the formula: 𝑭_𝐸𝐶𝑀_ = 𝜅_𝐸𝐶𝑀_ (𝒓_*b*𝑒*a*𝑑_ − 𝒓_𝐸𝐶𝑀_). The spring constant in this harmonic force 𝜅_𝐸𝐶𝑀_ is chosen larger than that of the spring connecting the blue bead to its neighbor, so that the contraction of the spring between the leading bead and the ECM particle, due to much lower mobility of the ECM particle, pulls the blue bead forward. Then two things can happen. When the blue and magenta particles touch each other, a new random ECM particle is selected for the next cycle of traction force. In case of when due to the movement of the blue bead, the spring connecting it to its preceding yellow bead extends to the length of 𝑙_0_, the weak spring is substituted by the strong spring with 𝜅_𝑠_, a new bead is added, and the process repeats. In our model, the traction force mechanism of neurite extension is the major mechanism of growth, with linear growth playing only a minor role.

Now that we described all the forces and growth mechanism acting in the model, we summarize the equations of motion that are then iterated numerically. As our simulations are pertinent to microscopic movements of growing neurites, it is justified to use the overdamped versions of the Newton’s law, where we can neglect the inertial terms in comparison to friction and other forces. In this case, the disbalance of the forces acting on a particle (bead) is dissipated via friction. Casting those force balance equations in the discretized version of the Euler approximation with the time increment 𝑑𝑡, we arrive at the following simple governing equations. For every *i*-th bead in the neurite the equations are: 𝛥𝒓_𝑛,𝑖_ = (𝑭_𝑠,𝑖−1_ + 𝑭_𝑠,𝑖+1_ + ∑_*j*_ 𝑭_𝑒𝑥𝑐,𝑖*j*_) ⋅ 𝜈_𝑛_ ⋅ 𝑑𝑡. For the ECM particles: 𝛥𝒓_𝐸𝐶𝑀,j_ = (𝑭_𝐸𝐶𝑀_ + ∑_*j*_ 𝑭_𝑒𝑥𝑐,𝑖j_) ⋅ 𝜈_𝐸𝐶𝑀_ ⋅ 𝑑𝑡. Where 𝑭_𝐸𝐶𝑀_ only acts on the ECM particle if it is bound to the leading bead of the neurite. Finally, for the leading bead the equation is: 𝛥𝒓_*b*𝑒*a*𝑑_ = (𝑭(𝑡)_𝑠,𝑡𝑟*a*𝑐_ + 𝑭_𝐸𝐶𝑀_ + ∑_*j*_ 𝑭_𝑒𝑥𝑐,𝑖*j*_)𝜈_*b*_ ⋅ 𝑑𝑡, where 𝑭_𝑠,𝑡𝑟*a*𝑐_(𝑡) is the spring force due to the extension of the leading bead. Additionally, the leading bead mobility was chosen to be larger than for the rest of the neurite beads. To be able to efficiently simulate the system containing a large enough number of ECM particles and neurites, we introduced an auxiliary lattice on top of the positions of the actual particles. This lattice allowed us to update only the positions of the beads that are in the immediate vicinity of other displacing particles, thus significantly reducing the computational time.

### Strategy of simulations

In our model, a neurite is represented by the chain of beads. The typical bead-to-bead distance is our approximation of how finely we want to resolve the shape of the neurite. The sizes of the ECM particles were chosen in accordance with the typical sizes of the gel beads used in the experimental realization of ECM in relation to the typical dimension of the growing neurite. All quantities in our simulations, at this point, are dimensionless, in accordance with the phenomenological nature of the model. Our strategy thus is to use the simulations to reproduce observables of the wild-type data and then rescale the numerical results to match the experimental observations. From that point, however, the scaling of the simulations and experiments are fixed. In the next step, we modulate the parameters of the model to recapitulate the mutant data, without changing the scaling,

### Parameters and units of simulations

As mentioned above, all quantities in this equation are dimensionless, so generally the numerical values of the parameters for this system will be chosen to be around 1. Significant deviations from that are indicating that a certain specific effect was aimed at. For the spatial dimension, the radius of the matrix particle was set to 1. The time step was chosen such that with given mobilities simulations are stable and do not produce oscillations in the movements of particles characteristic to dense systems with excluded volume interactions.

### In-silico simulation of MID1 WT and MID1 KO neurite growth dynamics

Please note, that unlike in the experimental data, we used the actual neurite outline to calculate the length, while in the data it is the length of the track followed by the neurite tip. To a good approximation, these two quantities are not different in our simulations. We also remark that the tortuosity is the quantity defined as the length of the trajectory divided by the end-to-end distance, and the end-to-end distance of a random-walk-like trajectory can be close to zero. As a result, the fluctuations of this quantity are rather strong even if we simulate a relatively large number of growing neurites.

### Numerical implementation

The simulations code was written in Python and can be run on a desktop machine. Results of the simulations were visualized using Ovito^76^. The source of the code is available under the link: https://github.com/matharkrv/neurite_simulation

### Statistics and reproducibility

Data were statistically analyzed with Microsoft Excel or R using statistical tests indicated throughout the manuscript. No statistical methods were used to predetermine sample size. The investigators were not blinded to allocation and outcome analysis. The experiments were not randomized.

### Data availability

The scRNA-seq data for this study have benn deposited in the European Nucleotide Archive (ENA) at EMBL-EBI under accession number xxx. The data reporting the details of the individual tracks is deposited on Zenodo 10.5281/zenodo.15213837. The source code for the *in-silico* simulation of neurite growth is available at https://github.com/matharkrv/neurite_simulation. The data that support the findings of this study are available from the corresponding authors upon reasonable request.

## Additional information

**Correspondence and requests for materials** should be addressed to M.K. and S.F.

## Acknowledgements

We thank the NGS Core Unit of the University Clinic Erlangen for excellent service regarding sequencing of the scRNA-seq libraries and Margherita Alfonsetti for assistance with tissue culture. We also thank Chiara Spear for assistance with organoid cutting. This work was supported by grants from the German Research Foundation (M.K: KA3125/2-1; M.K., C.D.L.: GRK2162/2; V.Z., M.K., S.F.: project 460333672 CRC1540 EBM), the Schram foundation (T287/29577/2017) to M.K., the Bavarian State Ministry of Sciences, Research, and the Arts (ForInter; F.2-F2412.30/1/24) to C.D.L., M.K., S.F., the Interdisciplinary Center for Clinical Research (IZKF) at the University Hospital of the FAU Erlangen-Nürnberg to M.K. (Jochen-Kalden funding programme N7, S1) and S.F. (E32).

## Author contributions

F.F. generated all organoids and performed molecular and cellular phenotyping, including the live imaging experiments for neurite outgrowth as well as tracking of neurites. Sa.F. together with F.F. processed the organoids for scRNA-seq. A.S.M. performed the analysis of the scRNA-seq data with guidance from S.F.. P.D. and M. Kr. performed in-depth analyses of neurite tracks and modelling of neurite behavior with guidance from V.Z.. S.T. generated the genome edited SOX11 and the TCF4 PSC lines under supervision of C.D.L.. A.L.L. assisted with PSC lines. S.K. generated the genome edited MID1 iPSC lines under supervision of Su.S.. S.F. performed image analyses. M.K. and S.F. jointly generated figures, conceptualized and supervised the work, and obtained funding. V.Z., S.F., M.K. wrote the manuscript, with all authors contributing corrections and comments.

## Tables and their legends

**Table S1.** Numerical values of the parameters used in the simulations to reproduce MID1 WT and MID1 KO data sets.

|  |  |  |
| --- | --- | --- |
| Radius of neurite beads | $R_n$ | 0.56 |
| Mobility of neurite beads | $v_n$ | 0.05 |
| Mobility of leading neurite beads | $v_b$ | 1 |
| Link range | $R_{link}$ | 15 |
| Cone angle (wild type) | $\theta_{link}$ | $30^\circ$ |
| Cone angle (mutated) | $\theta_{link}$ | $75^\circ$ |
| Critical spring length | $l_0$ | 1.2 |
| Growth rate | $\alpha$ | 10 |
| Spring constant growing | $\kappa_g$ | 0.1 |
| Spring constant | $\kappa_s$ | 4 |
| Excluded volume forces spring constant | $\epsilon$ | 1 |
| ECM bead radius | $R_{ECM}$ | 1 |
| ECM bead mobility | $v_{ECM}$ | 0.1 |
| Simulation time | $time$ | 100 |
| Simulation time steps | $dt$ | 0.002 |
| Simulation box dimensions | $box\ dim$ | 400 x 400 |
| Organoid dimensions | $L_x, L_y$ | 5, 25 |

**Table S2.**
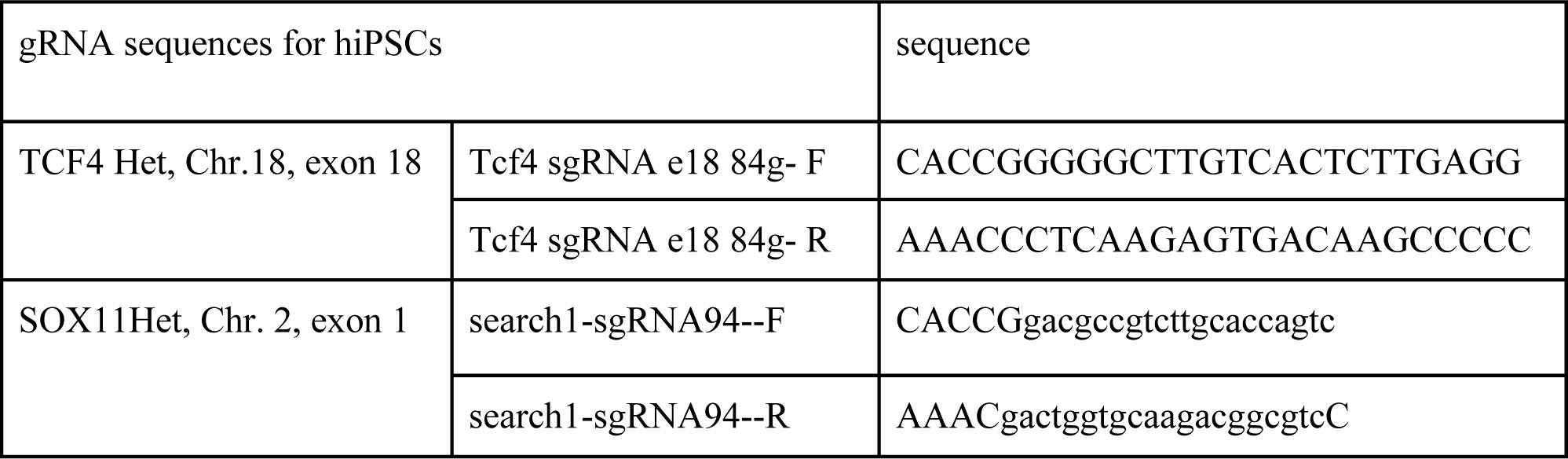
gRNAs used in this study.

## Supplementary Movies

**Movie M1**-example for neurite outgrowth from SOX11 WT organoid.

**Movie M2**-example for neurite outgrowth from SOX11 HET organoid.

**Movie M3**-example for neurite outgrowth from MID1 WT organoid.

**Movie M4**-example for neurite outgrowth from MID1 KO organoid.

**Movie M5**-example for neurite outgrowth from TCF WT organoid.

**Movie M6**-example for neurite outgrowth from TCF HET organoid.

**Movie M7**-numerical simulation of MID1 WT neurite outgrowth.

**Movie M8**-numerical simulation of MID1 KO neurite outgrowth.

## Supplementary Figures and Figure legends

**Figure S1.**
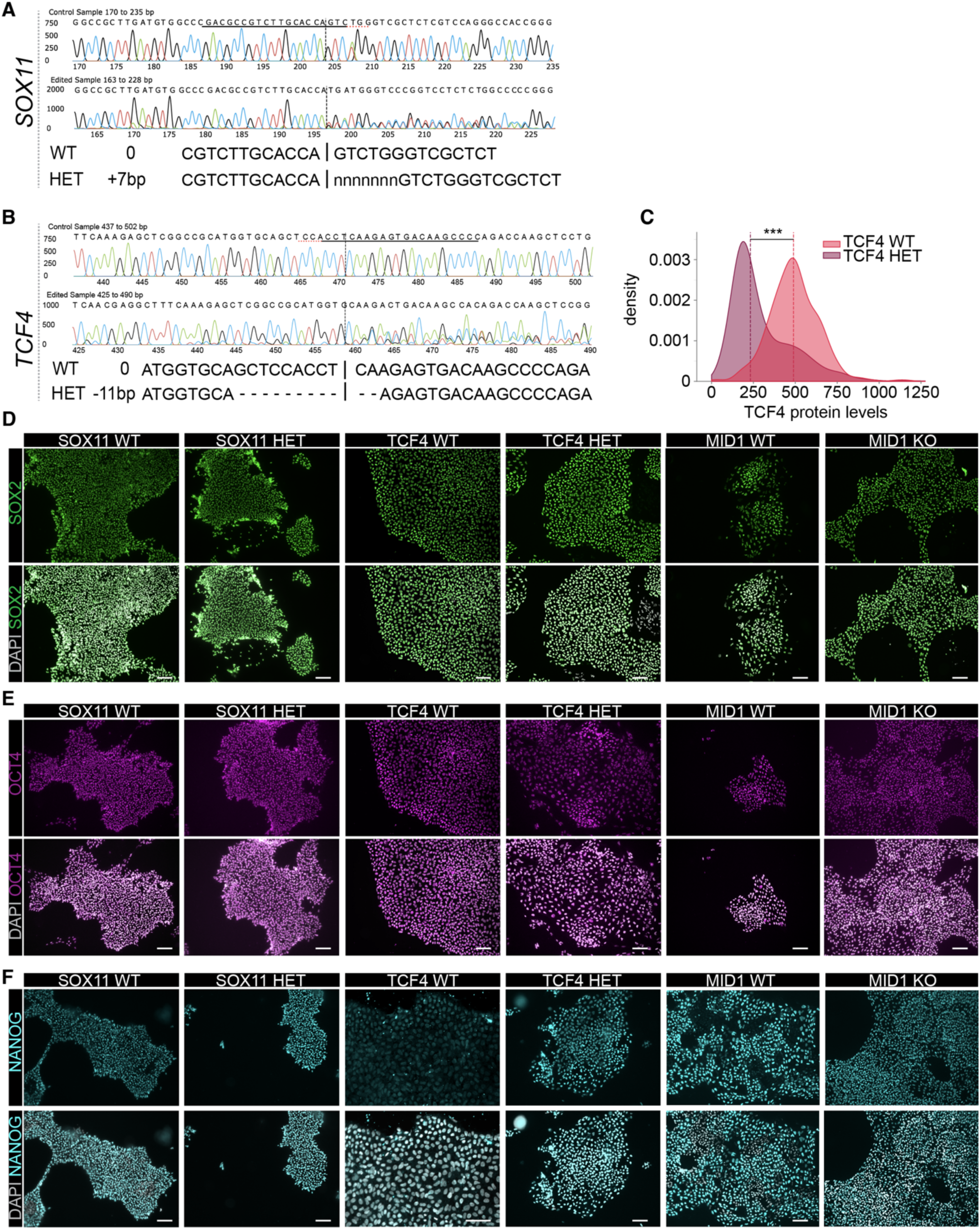
Characterization of PSC lines used in this study. A, B, ICE Analysis and validation of CRISPR editing in *SOX11* gene, +7 bp insertion, and *TCF4* gene, -11 bp insertion. **C,** Density distribution of TCF4 protein levels in TCF4 WT and TCF4 HET hPSC lines. Mann whitney U test. Exact P-value = 1.5 x10^-75^. **D, E, F** Representative immunofluorescence images of hPSC colonies from all lines used in this study, showing the expression of pluripotency markers SOX2 (green), OCT4 (magenta), and NANOG (blue), with nuclear staining using DAPI (gray). Scale bars =100µm.

**Figure S2.**
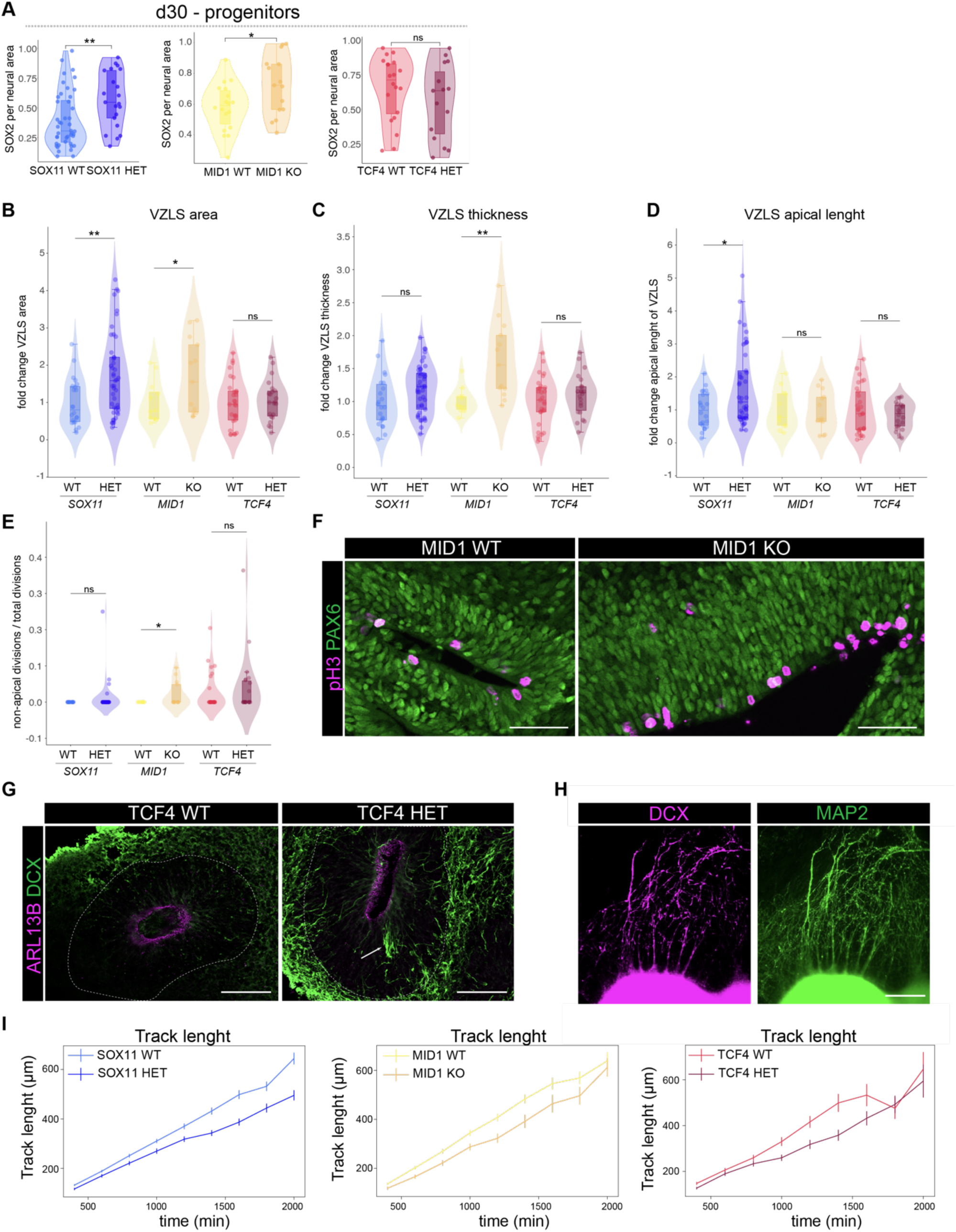
Transiently altered neurogenesis in SOX11, TCF4, and MID1 variant brain organoids. A, Violin, box, and jitter plots showing quantification of SOX2 positive pixels in neural areas (SOX2 and/or SOX2/MAP2) in whole organoid sections on d30. For SOX11 WT: *n* = 44; SOX11 HET: n = 21; exact *P* value = 0.0054. MID1 WT: *n* = 18; MID1 KO: n = 15, exact *P* value 0.024. TCF4 WT: *n* = 20; TCF4 HET: n = 15, exact *P* value 0.28. **B,** Violin, box, and jitter plots showing the quantification of the VZLS area revealing a significant increase in SOX11 HET and MID1 KO compared to their controls. SOX11 WT *n* = 22, SOX11 HET *n* = 45, MID1 WT *n* = 12, MID1 KO *n* = 13; TCF4 WT *n* = 29, TCF4 HET *n* = 22, exact *P* values (left to right): 0.0050, 0.0484, 0.7826. **C,** Quantification of thickness of the VZLS as shown by violin, box plots and jitter. SOX11 WT *n* = 22, SOX11 HET *n* = 45, MID1 WT *n* = 12, MID1 KO *n* = 13; TCF4 WT *n* = 29, TCF4 HET *n* = 22, exact *P* values (left to right): 0.0561, 0.0014, 0.3218. **D,** Violin, box plots and jitter showing the quantification of the apical length of VZLS as shown. SOX11 WT *n* = 22, SOX11 HET *n* = 45, MID1 WT *n* = 12, MID1 KO *n* = 13; TCF4 WT *n* = 29, TCF4 HET *n* = 22; exact *P* values (left to right): 0.0112, 0.8987, 0.3289. **E,** Violin, box plots and jitter showing the fraction of non-apical divisions per total divisions within VZLS across conditions. SOX11 WT *n* = 22, SOX11 HET *n* = 45, MID1 WT *n* = 12, MID1 KO *n* = 13; TCF4 WT *n* = 29, TCF4 HET *n* = 22; exact *P* values (left to right): 0.2937, 0.0459, 0.3992. For **A, B, C, D, E,** Individual dots represent single organoids. \*\*\**P*<0.001, \*\**P*<0.01, \**P*<0.5, ns = non-significant; dots represent individual organoids; one-way anova with tukey posthoc test was performed. **F,** Images showing pH3 (magenta) and PAX6 (green) in MID1 WT (left) and MID1 KO (right) organoids on d30. Note the presence of non-apically dividing cells in the MID1 KO condition. Scale bars = 100 µm. **G,** Images showing the cilia marker ARL13b (magenta) and the neuroblast marker DCX (green) in TCF4 WT (left) and TCF4 HET (right) organoids on d30. The white arrow points to mislocalized cells with DCX expression within the VZLS of TCF4 Het. Scale bars = 100 µm. **H**, Representative images showing neurite outgrowth from d60 brain organoid slices stained for the neuronal cytoskeleton markers DCX (magenta) and MAP2 (green). Scale bar = 200 µm. **I,** Line plot showing the track length (in µm) as a function of time (in min) for the SOX11 (left panel), the MID1 (middle panel), and the TCF4 conditions. Error bars indicate standard error of mean.

**Figure S3.**
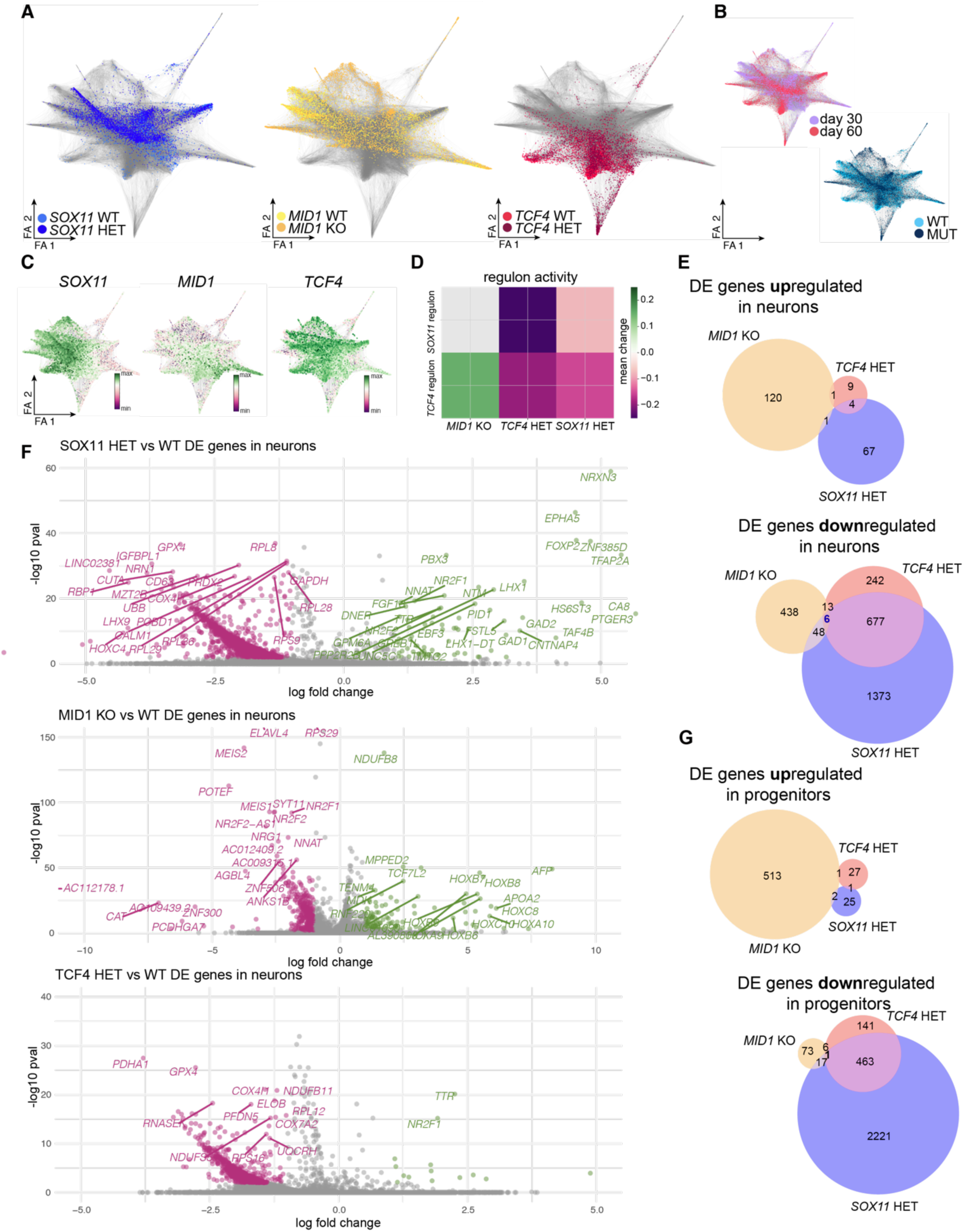
Molecular profiling of brain organoids unravels molecular nexuses. A, Force-directed graph embedding showing the transcriptomes of the cells of the individual conditions projected onto the embedding shown in Fig. 3A. Left panel shows SOX11 lines, middle panel shows MID1 lines, right panel shows TCF4 lines. **B,** Indication of the experimental days (d30, d60; upper panel) and the annotation as WT or MUT (= HET for SOX11 and TCF4, and KO for MID1) on the embedding shown in Fig. 3A. **C,** mRNA-expression of the *SOX11* (left), the *MID1* (middle), and the *TCF4* (right) gene projected onto embedding shown in Fig. 3A. For A, B, C, FA refers to force atlas. **D,** Heatmap indicating the mean change in the activity of the *SOX11* (upper row) and the *TCF4* regulon (lower row) in progenitors and neurons across experimental conditions. Gray color means no significant change. Wilcoxon test was used. **E,** Euler diagrams showing DE genes up- (upper panel) or downregulated (lower panel) in neurons across conditions. **F,** Volcano plots indicating the DE genes and the direction of deregulation across samples in neurons. **G,** Euler diagrams showing DE genes up- (upper panel) or downregulated (lower panel) in progenitors across conditions.

**Figure S4.**
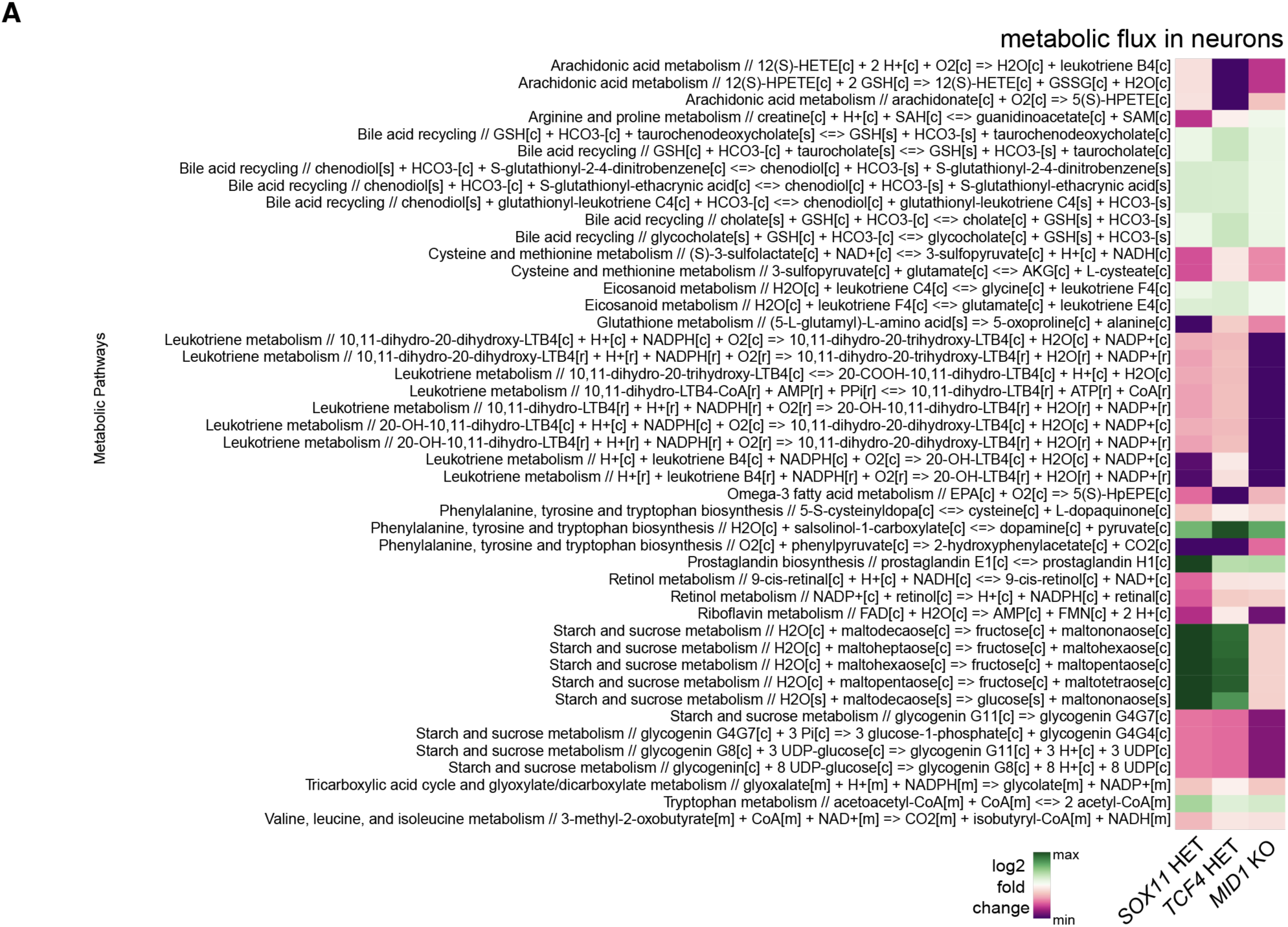
Molecular profiling of brain organoids suggests shared changes in metabolism. A, Heatmap showing extent of deregulation (log2 fold change) of significantly altered metabolic pathways including the indication of the exact biochemical reaction.

**Figure S5.**
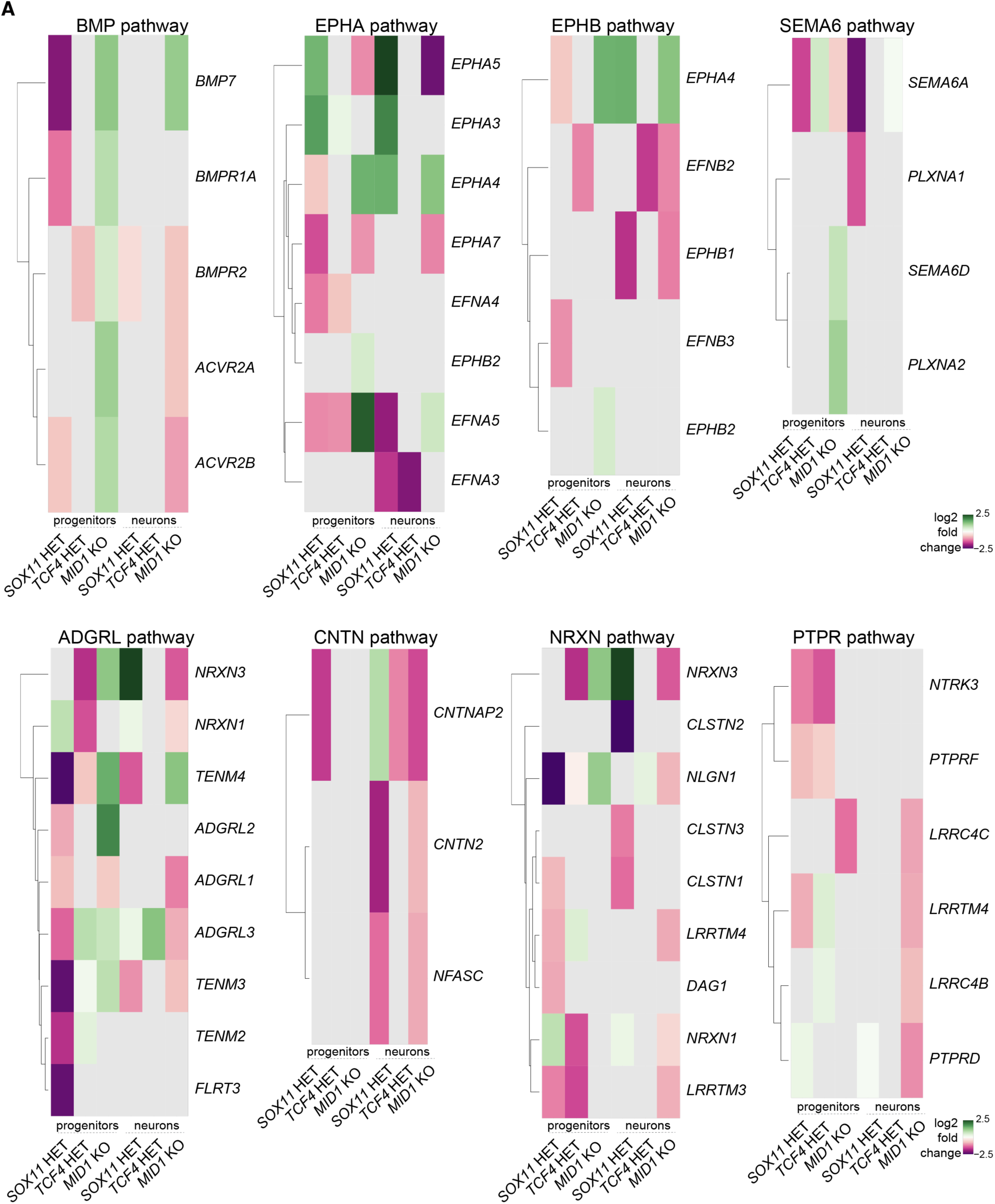
Assessment of cellular communication uncovers impaired signaling and reduced interactions with ECM. A, Heatmaps showing the expression of members of the 8 shared deregulated signaling pathways across conditions. Gray colors indicate non-significant changes.

